# Mucin-type O-glycosylation Landscapes of SARS-CoV-2 Spike Proteins

**DOI:** 10.1101/2020.07.29.227785

**Authors:** Yong Zhang, Wanjun Zhao, Yonghong Mao, Yaohui Chen, Jingqiang Zhu, Liqiang Hu, Meng Gong, Jingqiu Cheng, Hao Yang

## Abstract

The densely glycosylated spike (S) proteins that are highly exposed on the surface of severe acute respiratory syndrome coronavirus 2 (SARS-CoV-2) facilitate viral attachment, entry, and membrane fusion. We have previously reported all the 22 *N*-glycosites and site-specific *N*-glycans in the S protein protomer. Herein, we report the comprehensive and precise site-specific O-glycosylation landscapes of SARS-CoV-2 S proteins, which were characterized using high-resolution mass spectrometry. Following digestion using trypsin and trypsin/Glu-C, and de-N-glycosylation using PNGase F, we determined the mucin-type (GalNAc-type) O-glycosylation pattern of S proteins, including unambiguous *O*-glycosites and the 6 most common *O*-glycans occupying them, via Byonic identification and manual validation. Finally, 43 *O*-glycosites were identified in the insect cell-expressed S protein. Most glycosites were modified by non-sialylated *O*-glycans such as HexNAc(1) and HexNAc(1)Hex(1). In contrast, 30 *O*-glycosites were identified in the human cell-expressed S protein S1 subunit. Most glycosites were modified by sialylated *O*-glycans such as HexNAc(1)Hex(1)NeuAc(1) and HexNAc(1)Hex(1)NeuAc(2). Our results are the first to reveal that the SARS-CoV-2 S protein is a mucin-type glycoprotein; clustered *O*-glycans often occur in the N- and the C-termini of the S protein, and the *O*-glycosite and *O*-glycan compositions vary with the host cell type. These site-specific O-glycosylation landscapes of the SARS-CoV-2 S protein are expected to provide novel insights into the viral binding mechanism and present a strategy for the development of vaccines and targeted drugs.

## 1. INTRODUCTION

The spike (S) protein of severe acute respiratory syndrome coronavirus 2 (SARS-CoV-2) is an extensively N-glycosylated protein^1^ that protrudes from the virus surface and binds to the angiotensin-converting enzyme 2 (ACE2) receptor on host cells to mediate cell entry^2^. All 22 *N*-glycosites and *N*-glycans attached to asparagine (Asp, N) in a recombinant S protein protomer expressed in human and insect cells have been identified using high-resolution liquid chromatography–tandem mass spectrometry (LC-MS/MS)^3^. These *N*-glycosites are preferentially distributed in two functional subunits responsible for receptor binding (S1 subunit) and membrane fusion (S2 subunit)^3^. Site-specific N-glycosylation analysis can provide valuable insights into the infection mechanism and present a strategy for the development of vaccines^4^.

Unlike N-glycosylation, mucin-type O-glycosylation is initiated by the α-glycosidic attachment of *N*-acetylgalactosamine (GalNAc) to the hydroxyl group of serine (Ser, S) or threonine (Thr, T), which contains eight types of core structures (Core-1 to Core-8 *O*-glycans), and is involved in a variety of biological functions, such as the mediation of pathogenic binding to human receptors^4,5^. Moreover, O-glycosylation can influence proteolysis during antigen processing, which could prevent the formation of glycopeptides for further presentation to major histocompatibility complex (MHC) and the elicitation of immune response^6^. The S protein *O*-glycosites of SARS-CoV-2 have been predicted using computational analysis^7^, and Shajahan et al. (2020) identified two *O*-glycosites (T323 and S325) using LC-MS/MS^8^. However, mucin-type O-glycosylation often occurs in a cluster^9^. Hence, we believe that there are many mucin-type *O*-glycosites that have not been discovered as deciphering protein O-glycosylation remains a big challenge. The comprehensive and precise site-specific O-glycosylation analysis cannot be performed without appropriate sample preprocessing, analysis methods, and software^10–15^.

In the present study, we characterized the site-specific O-glycosylation of recombinant SARS-CoV-2 S proteins expressed in human and insect cells, using LC-MS/MS. Based on a complementary enzyme digestion strategy, we identified large-scale *O*-glycosites and their corresponding *O*-glycans in the recombinant S proteins, by extensive manual interpretation. The glycosite-specific occupancy by different glycoforms of S protein S1 subunits expressed in human and insect cells was resolved and compared. Detailed O-glycosylation profiles of S proteins are complementary to the N-glycosylation profiles and may help in the development of vaccines and therapeutic drugs.

## 2. EXPERIMENTAL SECTION

### 2.1. Materials and chemicals

Dithiothreitol (DTT), iodoacetamide (IAA), formic acid (FA), trifluoroacetic acid (TFA), Tris base, and urea were purchased from Sigma (St. Louis, MO, USA). Acetonitrile (ACN) was purchased from Merck (Darmstadt, Germany). Zwitterionic hydrophilic interaction liquid chromatography (ZIC-HILIC) materials were purchased from Fresh Bioscience (Shanghai, China). Recombinant SARS-CoV-2 S protein (S1+S2 ECD, His tag) expressed by insect cells (High Five) via a baculovirus, and S protein (S1, His tag) expressed by human embryonic kidney (HEK293) cells were purchased from Sino Biological (Beijing, China). Sequencing-grade trypsin and Glu-C were obtained from Enzyme & Spectrum (Beijing, China). A quantitative colorimetric peptide assay kit was purchased from Thermo Fisher Scientific (Waltham, MA, USA). Deionized water was prepared using a Milli-Q system (Millipore, Bedford, MA, USA). All other chemicals and reagents of the best available grade were purchased from Sigma-Aldrich or Thermo Fisher Scientific.

### 2.2. Protein digestion

Recombinant S proteins were proteolyzed using an in-solution protease digestion protocol. In brief, 50 μg of protein in a tube was denatured for 10 min at 95 °C. After reduction by DTT (20 mM) for 45 min at 56 °C and alkylation with IAA (50 mM) for 1 h at 25 °C in the dark, 2 μg of protease (trypsin or trypsin/Glu-C) was added to the tube and incubated for 16 h at 37 °C. After desalting using a pipette tip packed with a C18 membrane, the peptide concentration was determined using a peptide assay kit, based on the absorbance measured at 480 nm. The peptide mixtures were freeze-dried for further analysis.

### 2.3. Enrichment of intact glycopeptides and N-glycan removal

Intact *N*- and *O*-glycopeptides were enriched with ZIC-HILIC materials. Specifically, 20 μg of peptides was suspended in 100 μL of 80% ACN/0.2% TFA solution, and 2 mg of processed ZIC-HILIC materials was added to the peptide solution and incubated for 2 h at 37 °C. Finally, the mixture was transferred to a 200 μL pipette tip packed with a C8 membrane, and washed twice with 80% ACN/0.2% TFA. After enrichment, intact glycopeptides were eluted thrice with 70 μL of 0.1% TFA, and dried using a SpeedVac concentrator. The enriched intact glycopeptides were digested using 1 U PNGase F dissolved in 50 μL of 50 mM NH_4_HCO_3_ for 2 h at 37 °C. The reaction was terminated by adding 0.1% FA. The de-*N*-glycopeptides and *O*-glycopeptides were dried using a SpeedVac concentrator for further analysis.

### 2.4. Liquid chromatography-tandem mass spectrometry analysis

All the samples were analyzed using higher-energy collisional dissociation (HCD) in mass spectrometry (Orbitrap Fusion Lumos mass spectrometer). In brief, intact *O*-glycopeptides and de-*N*-glycopeptides were dissolved in 0.1% FA and separated on a column (ReproSil-Pur C18-AQ, 1.9 μm, 75 μm inner diameter, 20 cm length; Dr Maisch) over a 78 min gradient (buffer A, 0.1% FA in water; buffer B, 0.1% FA in 80% ACN) at a flow rate of 300 nL/min. MS1 was analyzed with a scan range (m/z) of 350–1550 at an Orbitrap resolution of 120,000. The RF lens, AGC target, maximum injection time, and exclusion duration were 30%, 1.0e6, 50 ms, and 15 s, respectively. MS2 was analyzed with an isolation window (m/z) of 2 at an Orbitrap resolution of 15,000. The AGC target, maximum injection time, and HCD type were 5.0e4, 80 ms, and 35%, respectively.

### 2.5. Data analysis

Raw data files were searched against the SARS-CoV-2 S protein sequence using Byonic™ software (version 3.6.0, Protein Metrics, Inc.)^16^, with the mass tolerance for precursors and fragment ions set at ±10 and ±20 ppm, respectively. Two missed cleavage sites were subjected to trypsin or trypsin/Glu-C digestion. The fixed modification was carbamidomethyl (C), and the variable modifications included oxidation (M), acetyl (protein N-term), and de-amidation (N). In addition, the 6 most common mucin-type *O*-glycans (HexNAc(1) with mass of 203.079 Da; HexNAc(2) with mass of 406.159 Da; HexNAc(1)Hex(1) with mass of 365.132 Da; HexNAc(2)Hex(1) with mass of 568.212 Da; HexNAc(1)Hex(1)NeuAc(1) with mass of 656.228 Da; and HexNAc(1)Hex(1)NeuAc(2) with mass of 947.323 Da) were specified as *O*-glycan modifications for intact *O*-glycopeptides. We then checked the protein database options, including the decoy database. All other parameters were set to the default values, and protein groups were filtered using a 1% false discovery rate, based on the number of hits obtained for the searches against the databases. Stricter quality control methods for intact *O*-glycopeptide identification were implemented; they required a score of not less than 300, and at least 6 amino acids to be identified. Furthermore, all the glycopeptide-spectrum matches (GPSMs) were examined manually and filtered using the following standard criteria: a. GPSMs were accepted if there were at least two glycan oxonium ions and at least 6 b/y ions in the peptide backbone. b. The unambiguously identified *O*-glycosites were not to be hindered by the surrounding potential *O*-glycosites (Ser or Thr). In addition, the high-confidence *O*-glycosites had to be identified repeatedly at least twice. *O*-glycosite conservation analysis was performed using R software packages. Model building based on the Cryo-EM structure (PDB: 6VSB) of the SARS-CoV-2 S protein was performed using PyMOL.

## 3. RESULTS AND DISCUSSION

### 3.1. Strategy for intact O-glycopeptide analysis

Our previous study revealed site-specific N-glycosylation of recombinant S proteins^3^. However, comprehensive and precise site-specific O-glycosylation analysis of the SARS-CoV-2 S protein has not been performed. In the present study, we aimed to characterize the site-specific mucin-type O-glycosylation landscapes of SARS-CoV-2 recombinant S proteins.

The strategy for intact *O*-glycopeptide analysis is shown in Figure 1A. The recombinant SARS-CoV-2 S proteins were digested using trypsin or a mixture of trypsin and Glu-C to cover as many potential *O*-glycosites as possible. Then, intact glycopeptides were enriched using ZIC-HILIC^17^, and de-N-glycosylated with PNGase F to avoid interference from non-glycopeptides and *N*-glycopeptides. Finally, intact *O*-glycopeptides were analyzed using a high-resolution mass spectrometer, and their mass spectra were characterized using Byonic™ and validated manually^18^. The S protein expressed in insect cells contained 1209 amino acids (residues 16–1,213) and included 94 Thr and 92 Ser residues as potential mucin-type *O*-glycosites. The spike protein S1 subunit expressed in human cells contained 681 amino acids (residues 16–685) and included 57 Thr and 50 Ser residues as potential mucin-type *O*-glycosites (Figure S1). Combined digestion can cover more potential sites, including the reported and predicted *O*-glycosites^19^. As shown in Figure 1B, there was strong mass spectra evidence for the presence of O-glycosylation at site Thr323, as reported recently^8^. The representative HCD-MS/MS spectra of intact *O*-glycopeptide ^320^VQPTESIVR^328^ with GalNAcGal on site Thr323 (Figure 1B) indicated sufficient fragment ions (b ions, y ions, and oxonium ions) to confirm the presence of O-glycosylation on the target site of the peptide. In addition, the experimental data and manual interpretation showed that intact *O*-glycopeptide ^320^VQPTESIVR^328^ could be O-glycosylated on a single site, i.e., Ser325 (Figure 1C), or both sites, i.e., Thr323 and Ser325 (Figure 1D). These results showed that our strategy was feasible for mucin-type O-glycosylation profiling.

**Figure 1.**
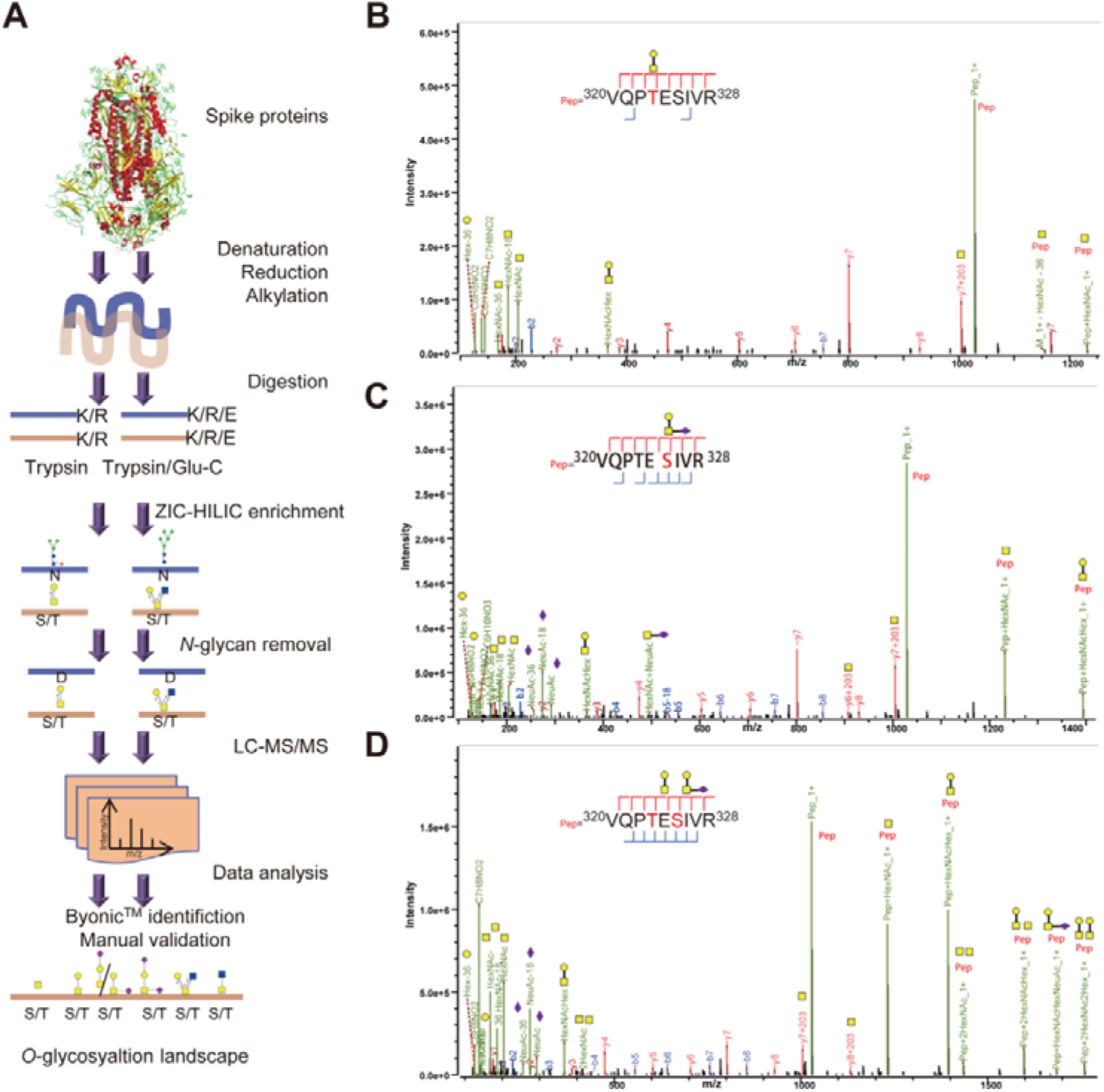
Site-specific *O*-glycosylation profiling of SARS-CoV-2 spike proteins. **A.** SARS-CoV-2 spike proteins expressed in insect or human cells were digested using trypsin or a mixture of trypsin and Glu-C. After ZIC-HILIC enrichment and PNGase F digestion, intact *O*-glycopeptides were analyzed using a high-resolution mass spectrometer, and their spectra were characterized using Byonic™ software and validated manually. **B.** SCE-HCD-MS/MS spectrum of reported representative *O*-glycopeptide ^320^VQPTESIVR^328^ with deduced GalNAcGal glycan detected on site Thr323 of human spike protein subunit 1. **C.** SCE-HCD-MS/MS spectrum of this *O*-glycopeptide with deduced GalNAcGalNeuAc glycan detected on site Ser325. **D.** SCE-HCD-MS/MS spectrum of this *O*-glycopeptide with deduced GalNAcGal glycan on site Thr323 and deduced GalNAcGalNeuAc glycan on site Ser325.

### 3.2. Site-specific O-glycosylation profiling of recombinant SARS-CoV-2 S protein expressed in insect cells

The S protein produced by the baculovirus insect cell expression system contained 186 potential *N*-glycosites. Using our aforementioned strategy, a total of 43 *O*-glycosites were assigned unambiguously with high-quality spectral evidence (Table S1 and Figure S2). Forty *O*-glycosites, except S477, T572, and T732 could be identified repeatedly using trypsin alone. Moreover, 40 *O*-glycosites, except S325, T333, and T1066, could be identified repeatedly using trypsin combined with Glu-C (Figure 2A). Hence, although trypsin digestion can yield good identification results, trypsin combined with Glu-C digestion should be considered as complementary step. Furthermore, we mapped these high-confidence *O*-glycosites to the amino sequences, and found that the *O*-glycosites clustered in several areas, especially in the N- and C-termini of the S protein (Figure 2B). It is notable that the *O*-glycosites T323, S325, T333, S345, and S477 were located in the receptor-binding domain (RBD). This was the first result to reveal that the SARS-CoV-2 S protein is a mucin-type glycoprotein. In addition, the number of O-glycosylated Thr residues (25) was higher than that of O-glycosylated Ser residues (18) (Figure 2B). This result is consistent with those of previous studies on O-glycoproteomics^18^. Finally, a global analysis of site-specific O-glycosylation of the S protein was performed (Figure 2C). Six mucin-type *O*-glycan compositions were identified on these sites, including HexNAc(1), HexNAc(2), HexNAc(1)Hex(1), HexNAc(2)Hex(1), HexNAc(1)Hex(1)NeuAc(1), and HexNAc(1)Hex(1)NeuAc(2). Specifically, HexNAc(1)Hex(1), HexNAc(1), HexNAc(2)Hex(1), HexNAc(1)Hex(1)NeuAc(1), HexNAc(2), and HexNAc(1)Hex(1)NeuAc(2) compositions corresponded to 40, 30, 21, 18, 11, and 7 glycosites, respectively. Most glycosites contained at least two types of *O*-glycans, a majority of which were non-sialylated (Figure 2C). These results indicated the clustered mucin-type *O*-glycans on the recombinant SARS-CoV-2 S protein expressed in insect cells.

**Figure 2.**
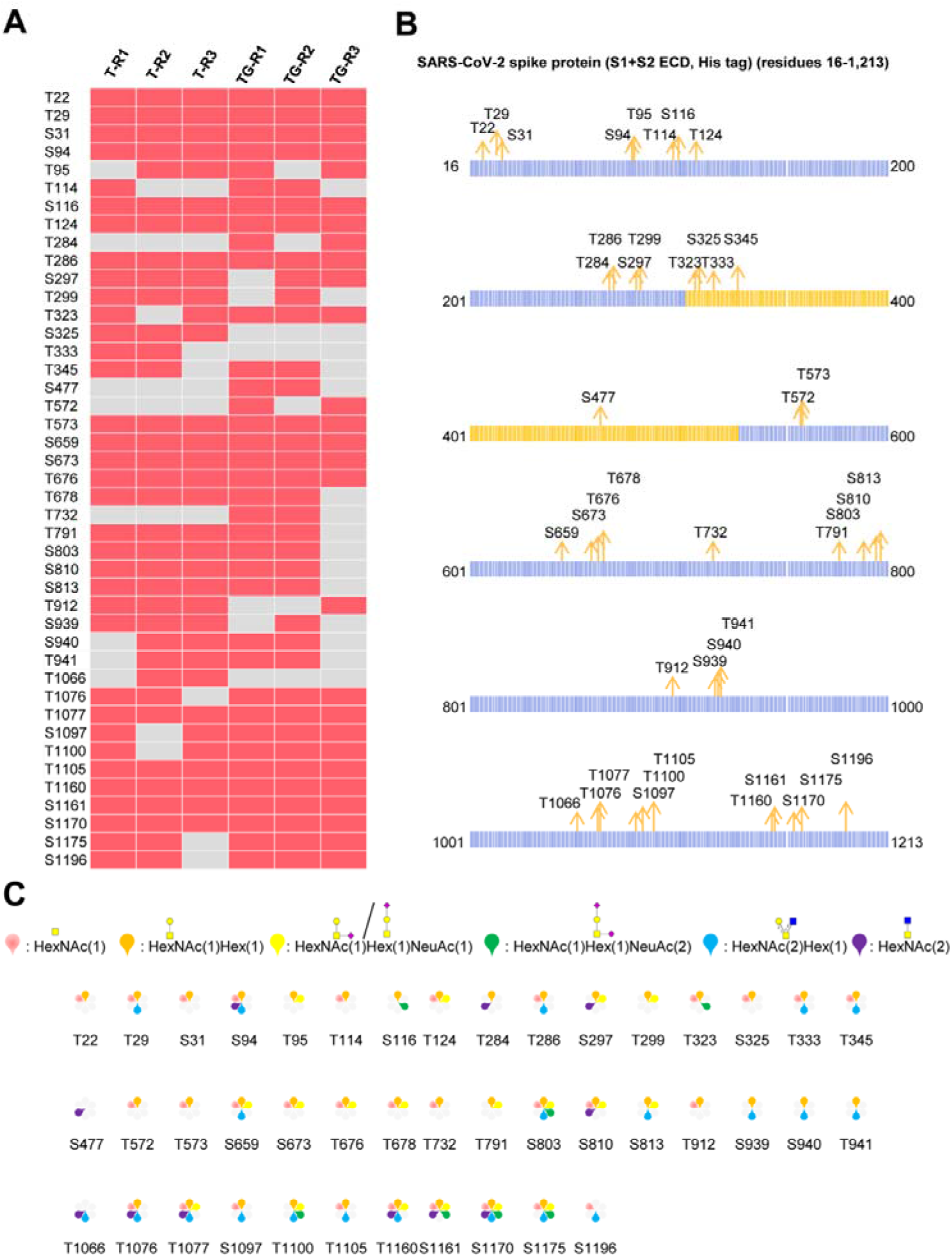
Site-specific *O*-glycosylation characterization of recombinant SARS-CoV-2 S protein (S1+S2 ECD, His tag) expressed in insect cells. **A.** High-confidence *O*-glycosites identified using trypsin (T) or typsin/Glu-C (TG) in three replicates. **B.** Mapping of identified *O*-glycosites to amino acid sequences. **C.** *O*-glycan compositions on each site.

### 3.3. Site-specific O-glycosylation profiling of recombinant SARS-CoV-2 S protein expressed in human cells

The recombinant SARS-CoV-2 S protein S1 subunit produced by the human cell expression system was used for analysis of the site-specific *O*-glycans, as the *O*-glycan compositions in insect cells could be different from those in human cells. Using our aforementioned strategy, 30 high-confidence *O*-glycosites (20 O-glycosylated Thr and 10 O-glycosylated Ser residues) were assigned unambiguously (Table S2 and Figure S3). Twenty-four and twenty-seven *O*-glycosites were identified repeatedly using trypsin and a mixture of trypsin/Glu-C, respectively (Figure 3A). The results showed that the two digestion methods were complementary for *O*-glycosite identification. Furthermore, we mapped these 30 *O*-glycosites to the amino sequences. We found that the *O*-glycosites mainly clustered at the N- and C-termini of the S1 subunit and RBD (Figures 3B and 3C). It is notable that two conserved *O*-glycosites, T323 and S325, were located in the RBD of the S1 subunit, and played a critical role in viral binding with hACE2 receptors^20,21^. A global analysis of site-specific O-glycosylation of the S1 subunit was performed. HexNAc(1)Hex(1)NeuAc(2), HexNAc(1)Hex(1), HexNAc(1)Hex(1)NeuAc(1), HexNAc(2), HexNAc(1), and HexNAc(2)Hex(1) compositions corresponded to 29, 25, 24, 16, 10, and 9 glycosites, respectively. Most glycosites contained at least two types of *O*-glycans, a majority of which were sialylated (Figure 3D). These results indicated the more complex mucin-type O-glycosylation and the heterogeneity of *O*-glycan compositions on the recombinant SARS-CoV-2 S protein expressed in human cells.

**Figure 3.**
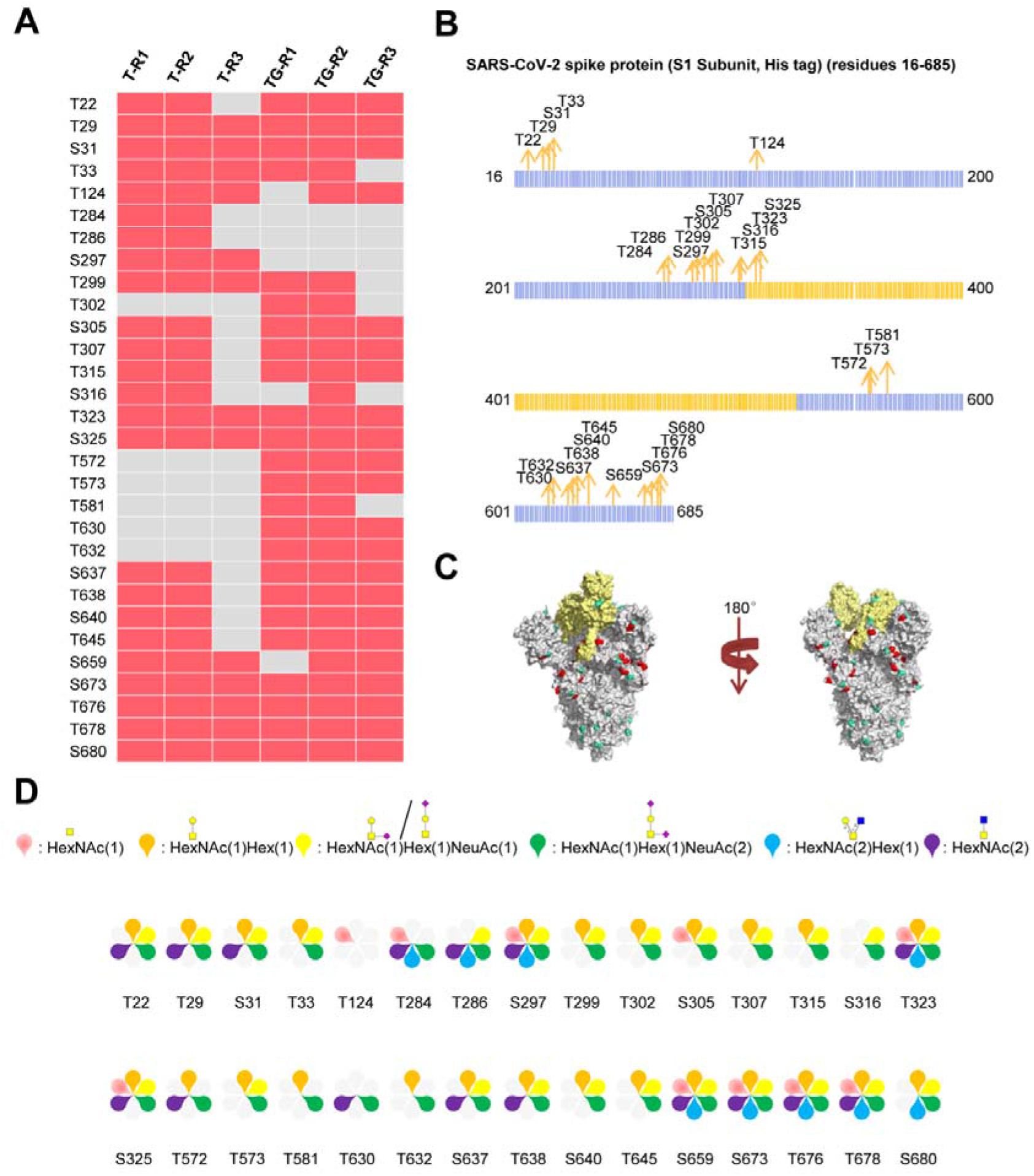
Site-specific *O*-glycosylation characterization of SARS-CoV-2 S protein (S1, His tag) expressed in human cells. **A.** *O*-glycosites identified using trypsin (T) or typsin/Glu-C (TG) in three replicates. **B.** Mapping of identified *O*-glycosites to amino acid sequences. **C.** *O*-glycosites (red) and *N*-glycosites (blue) in three-dimensional structure of SARS-CoV-2 S protein trimers (PDB code: 6VSB). **D.** *O*-glycan compositions on each site.

### 3.4. Comparison of O-glycosylation landscapes of S1 subunits expressed in insect and human cells

Based on the above findings, we further compared the O-glycosylation landscapes of the S1 subunits expressed in insect and human cells. Twenty-three *O*-glycosites were present in the S1 subunit expressed in insect cells (Figure 4A). In contrast, 30 *O*-glycosites were present in the S1 subunit expressed in human cells (Figure 4B). In addition, 16 conserved *O*-glycosites (T22, T29, S31, T124, T284, T286, S297, T299, T323, S325, T572, T573, S659, S673, T676, and T678) were identified in the S1 subunits expressed in insect and human cells, including two sites, T323 and S325, located in the RBD. Seven and fourteen unique *O*-glycosites were identified in the insect and human cell–produced S1 subunits, respectively (Figure 4C). Furthermore, the number of S1 subunit *O*-glycosites occupied by each type of *O*-glycan compositions was very different. Most *O*-glycosites of the insect cell–produced S1 subunit contained HexNAc(1)Hex(1) and HexNAc(1). On the other hand, most *O*-glycosites of the human cell–produced S1 subunit contained HexNAc(1)Hex(1)NeuAc(2), HexNAc(1)Hex(1)NeuAc(1), and HexNAc(1)Hex(1) (Figure 4D). These results implied that the *O*-glycosite and *O*-glycan compositions varied with the host cell type, which could be taken into account when using the recombinant proteins for vaccine and drug development.

**Figure 4.**
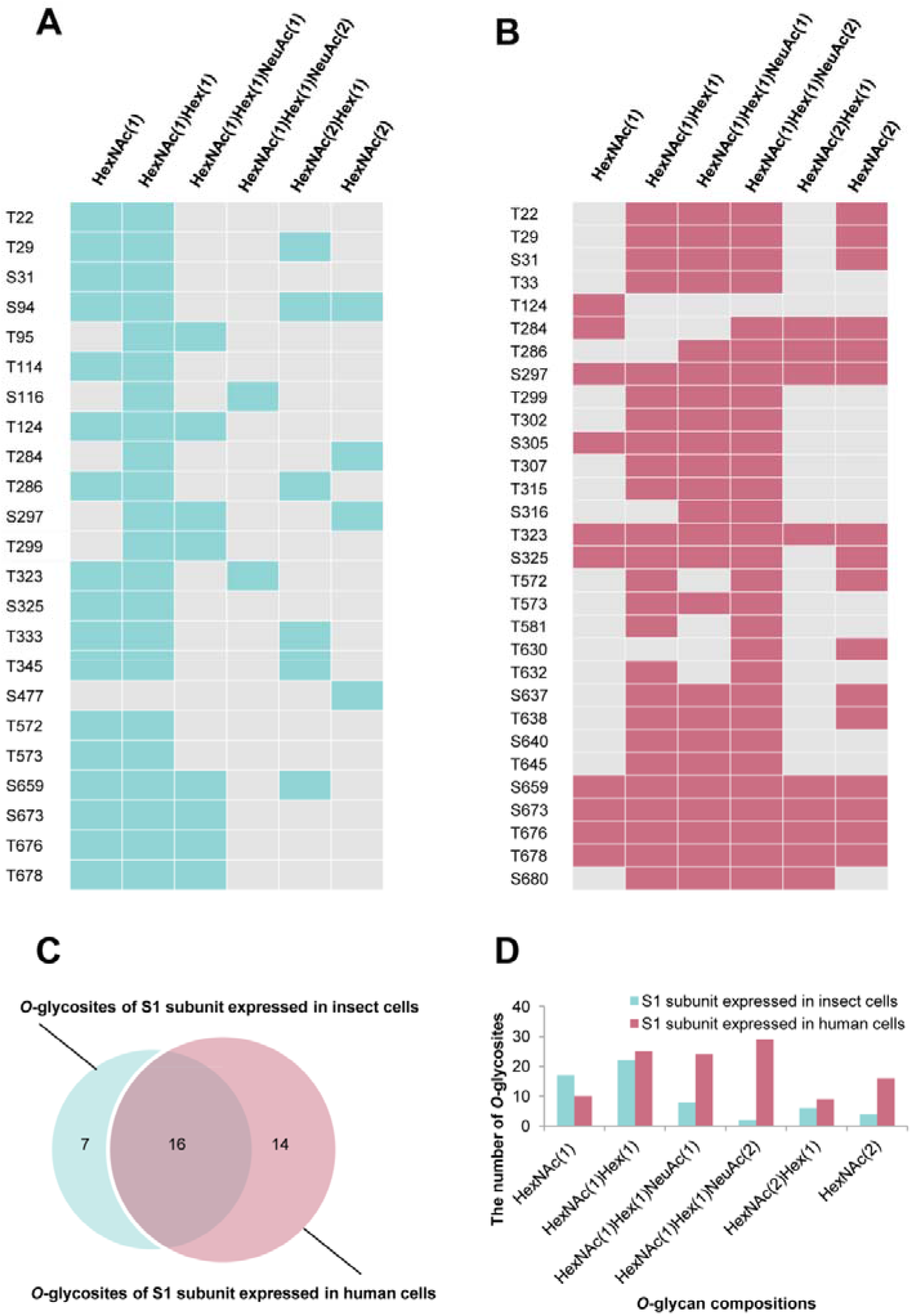
Comparison of site-specific *O*-glycosylation modifications of S1 subunits expressed in insect or human cells. **A.** *O*-glycan compositions in each glycosite of S1 subunit expressed in insect cells. **B.** *O*-glycan compositions in each glycosite of S1 subunit expressed in human cells. **C.** Comparison of *O*-glycosites of S1 subunits expressed in different expression systems. **D.** Number of S1 subunit *O*-glycosites attached by each type of *O*-glycan composition.

## 4. CONCLUSIONS

In this study, we profiled a comprehensive site-specific O-glycosylation pattern of SARS-CoV-2 S proteins using optimized experimental procedure and high resolution mass spectrometry. Forty-three *O*-glycosites were identified in insect cell–expressed S protein, and most of them were non-sialylated. In contrast, 30 *O*-glycosites were identified in human S protein, and most of them were sialylated. Our results revealed that the SARS-CoV-2 S protein was modified by clustered mucin-type *O*-glycans, and that the *O*-glycosite and *O*-glycan compositions varied with the host cell type.

## Supporting information

Table S1-Intact O-glycopeptides of SARS-CoV-2 spike protein expressed in insect cells

Table S2-Intact O-glycopeptides of SARS-CoV-2 spike protein expressed in human cells

## ACKNOWLEDGMENTS

This work was funded by grants from the National Natural Science Foundation of China (31901038), Department of Science and Technology of Sichuan Province (2020YFH0029), 1.3.5 Project for Disciplines of Excellence, West China Hospital, Sichuan University (ZYGD18014), and Chengdu Science and Technology Department Foundation (2020-YF05-00240-SN).

## Appendix A. Supplementary material

Supplementary data associated with this article can be found in the online version.

## Supporting Information

**Supplementary Figure S1.** Potential *O*-glycosites of SARS-CoV-2 S proteins expressed in insect and human cells

**Supplementary Figure S2.** Spectra of intact *O*-glycopeptides of SARS-CoV-2 S protein expressed in insect cells with ambiguously assigned *O*-glycosites

**Supplementary Figure S3.** Spectra of intact *O*-glycopeptides of SARS-CoV-2 S protein expressed in human cells with ambiguously assigned *O*-glycosites

