## Supporting_Information for "Mucin-type O-glycosylation Landscapes of SARS-CoV-2 Spike Proteins"

**Supplementary Figures:**

**Supplementary Figure S1.** Potential *O*-glycosites of SARS-CoV-2 S proteins expressed in insect and human cells

**Supplementary Figure S2.** Spectra of intact *O*-glycopeptides of SARS-CoV-2 S protein expressed in insect cells with ambiguously assigned *O*-glycosites

**Supplementary Figure S3.** Spectra of intact *O*-glycopeptides of SARS-CoV-2 S protein expressed in human cells with ambiguously assigned *O*-glycosites

**Supplementary Figure S1.** Potential *O*-glycosites of SARS-CoV-2 S proteins expressed in insect and human cells

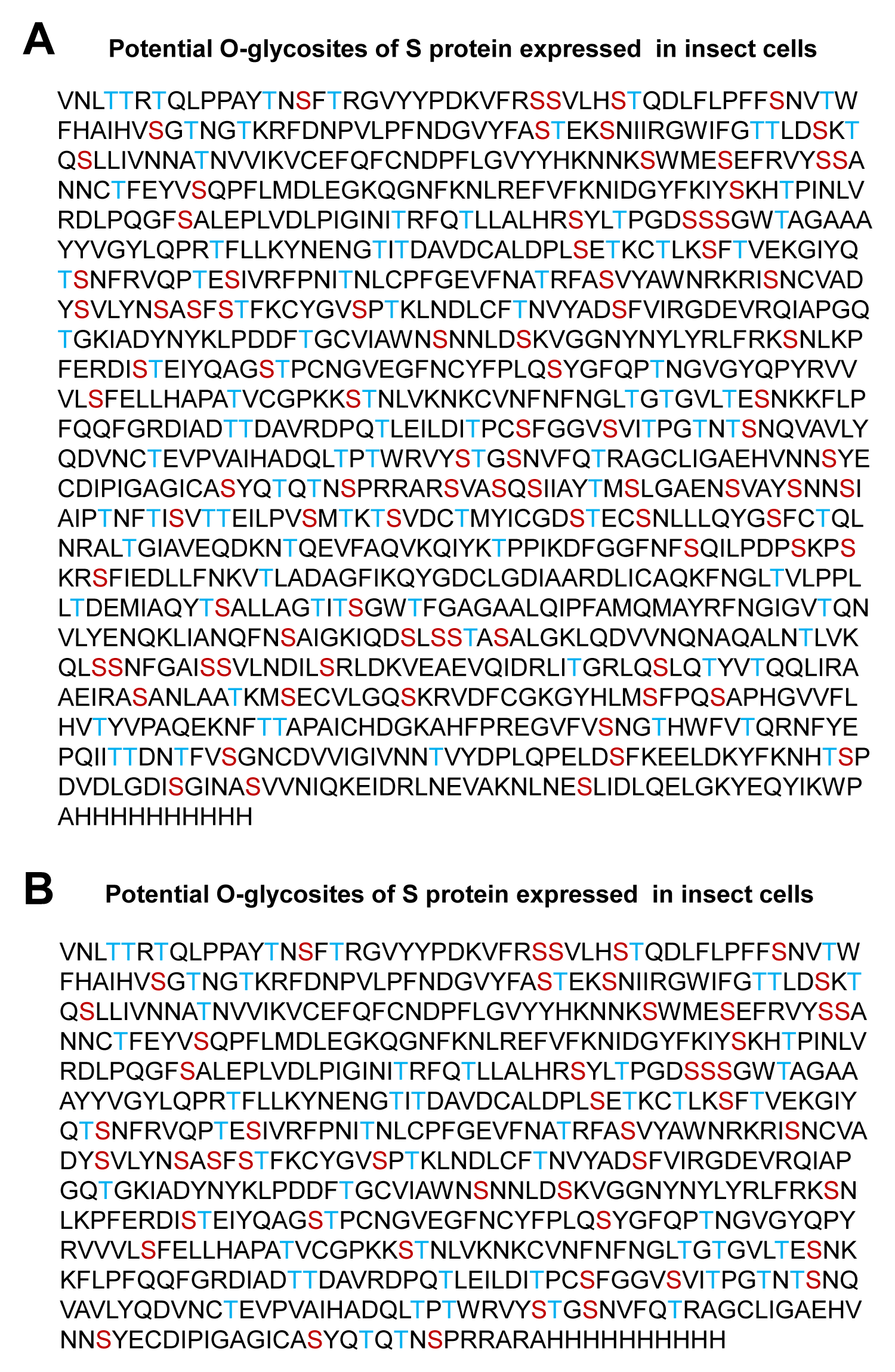

**Supplementary Figure S2.** Spectra of intact *O*-glycopeptides of SARS-CoV-2 S protein expressed in insect cells with ambiguously assigned *O*-glycosites

**T22 & S31**

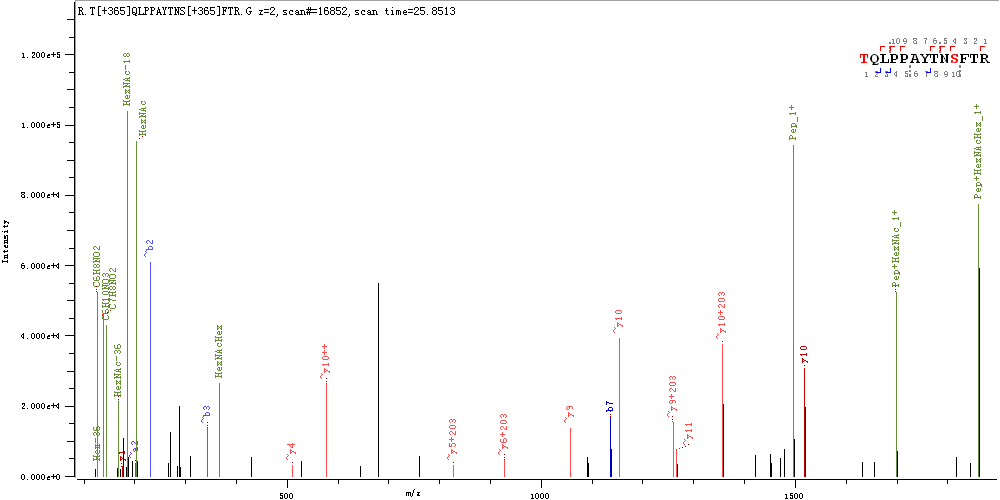

**T29**

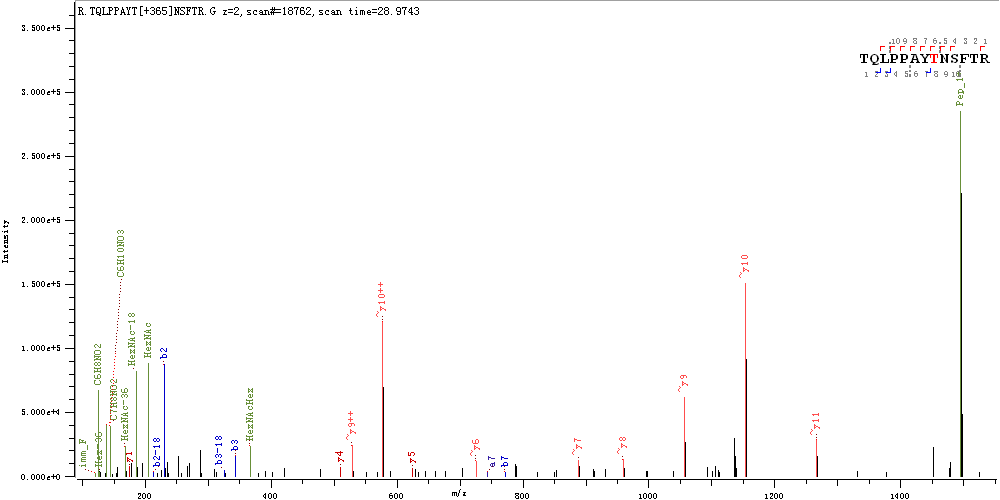

**S94**

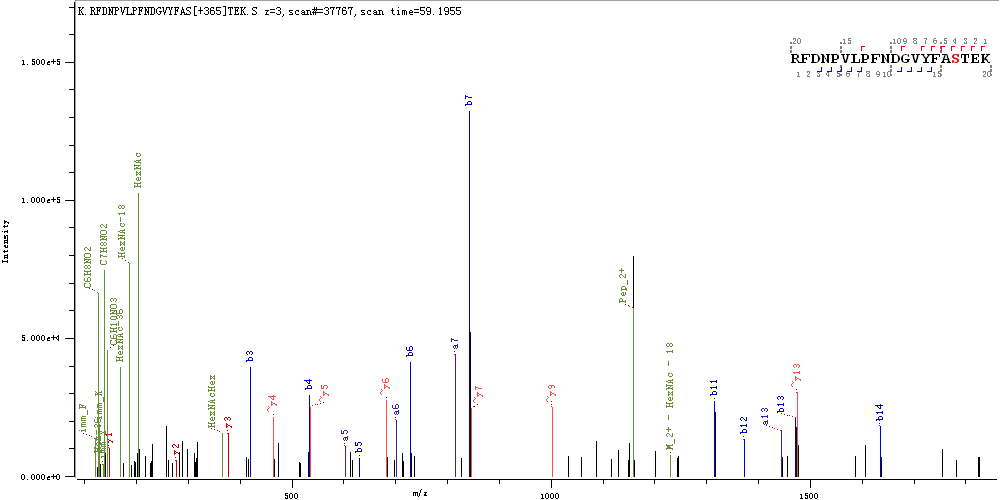

**S94 & T95**

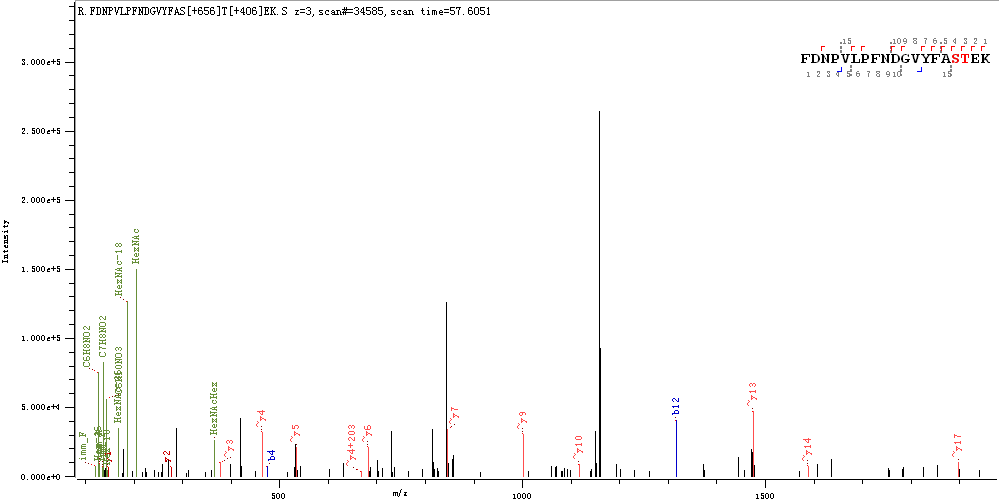

**T114 & S116 &T124**

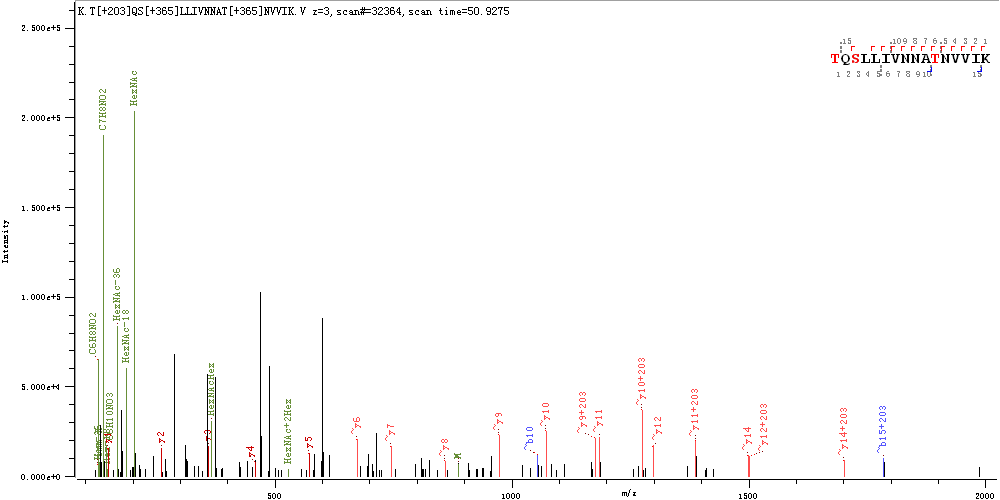

**T284 & T286 & S297**

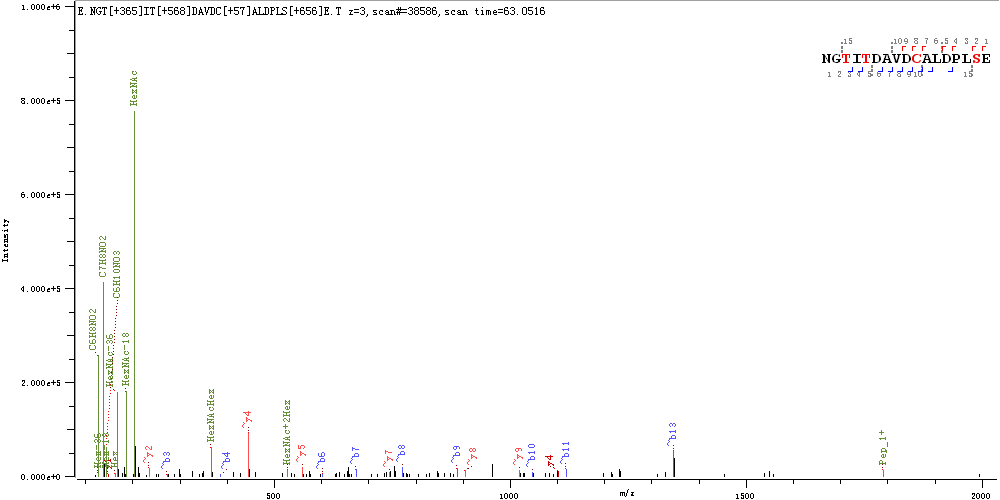

**T286 & S297&T299**

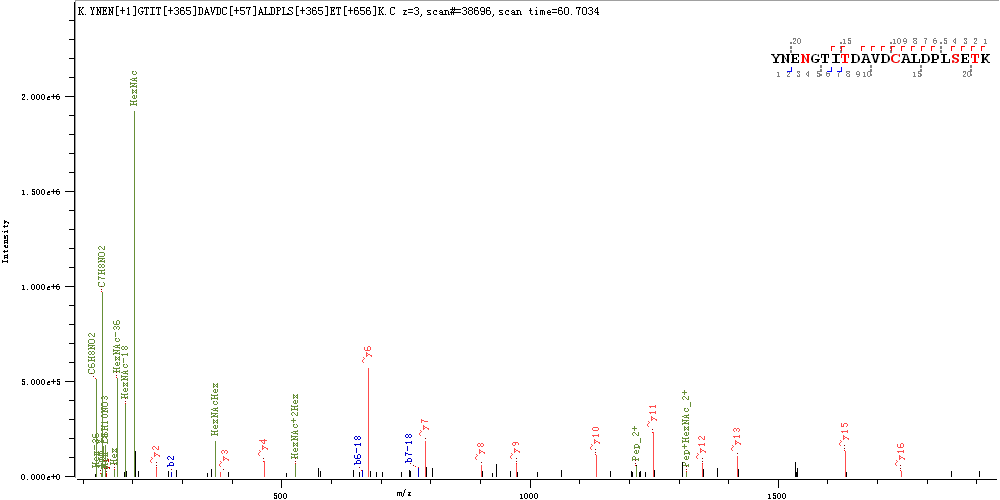

**T323**

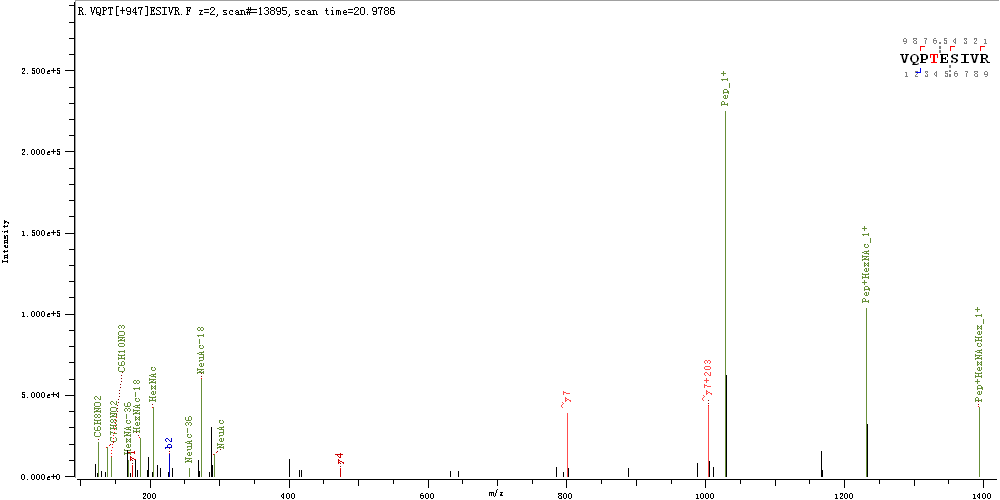

**S325**

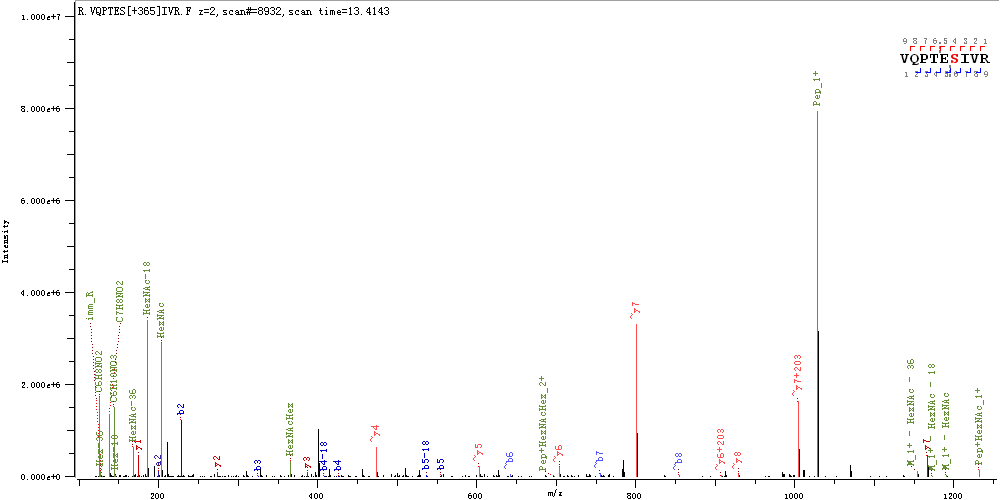

**T333 & T345**

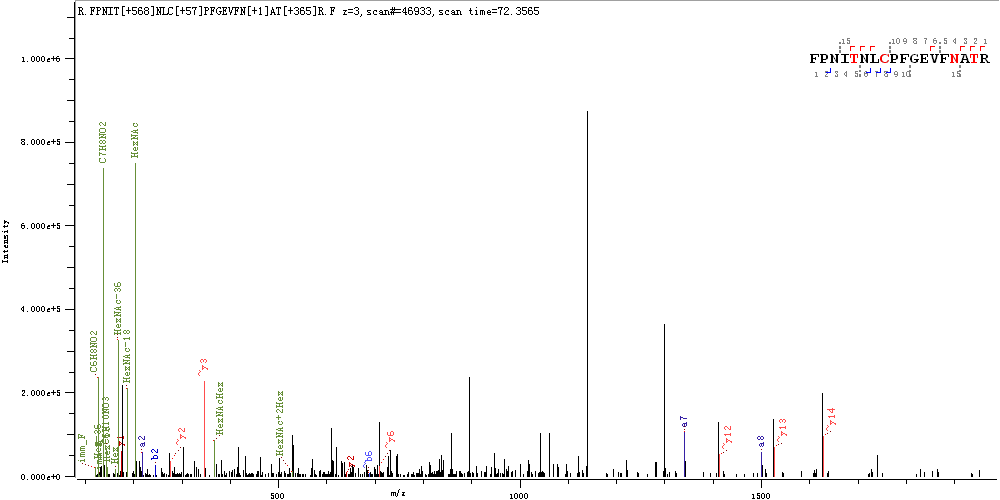

**S477**

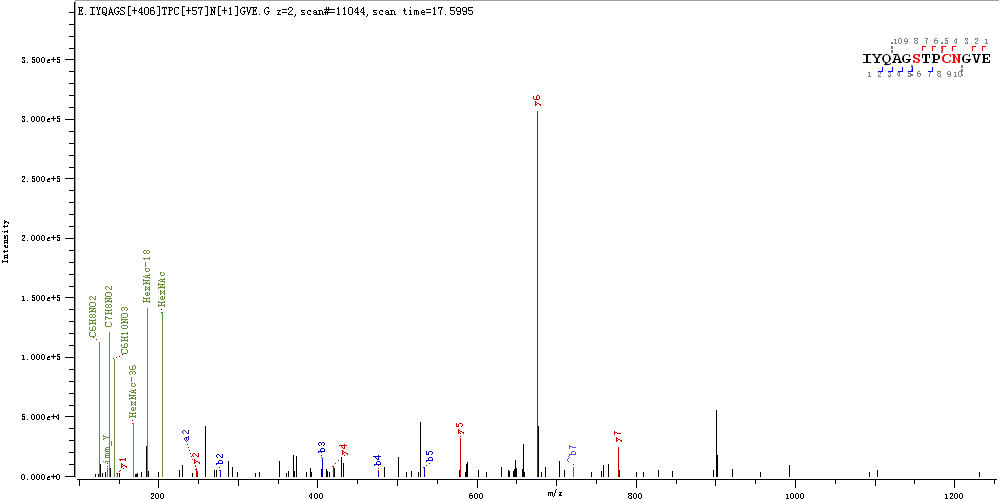

**T572**

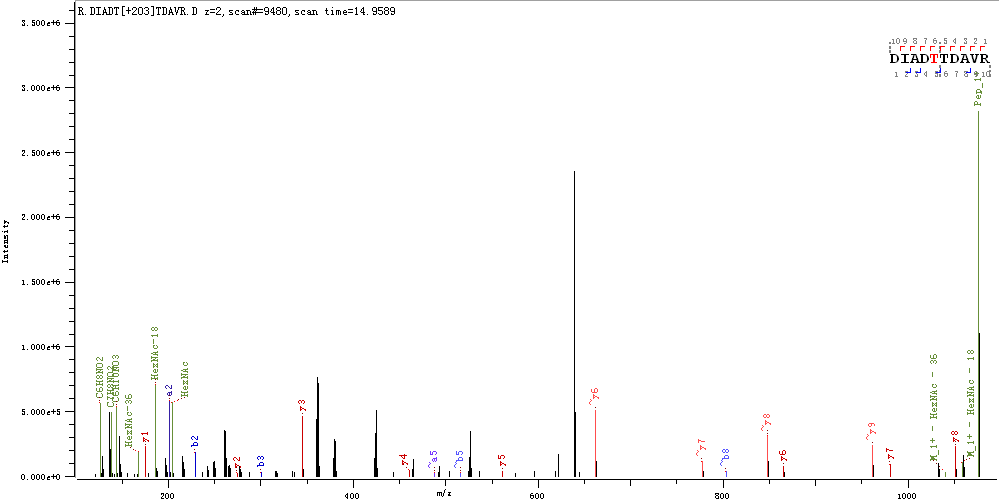

**T573**

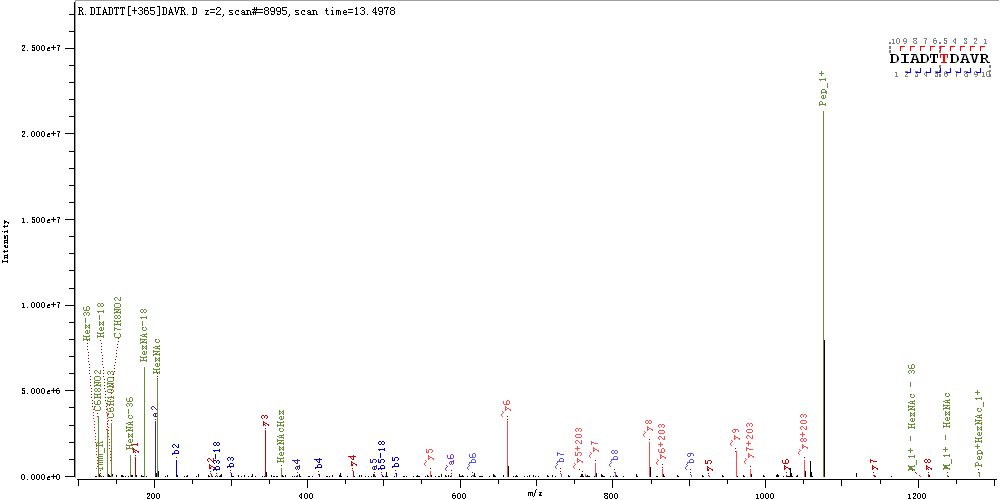

**S659 & T676 & T678**

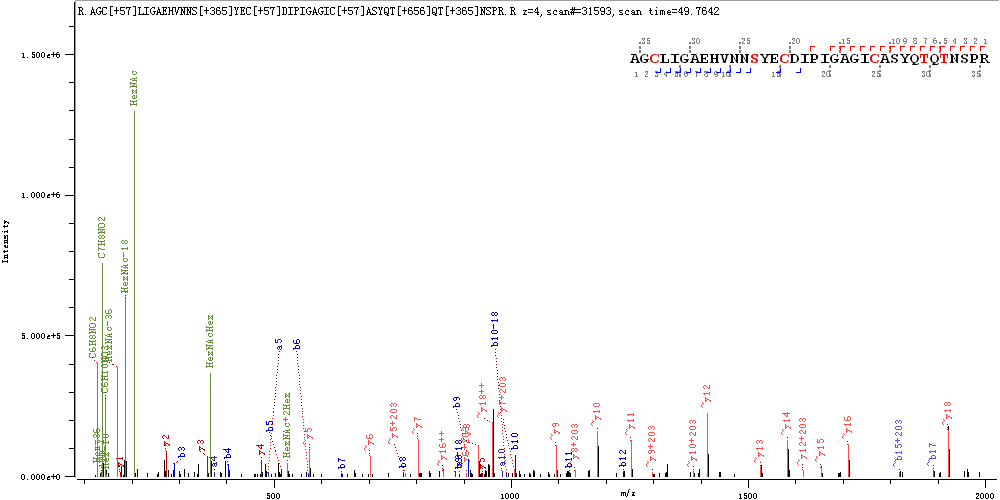

**S659 & S673**

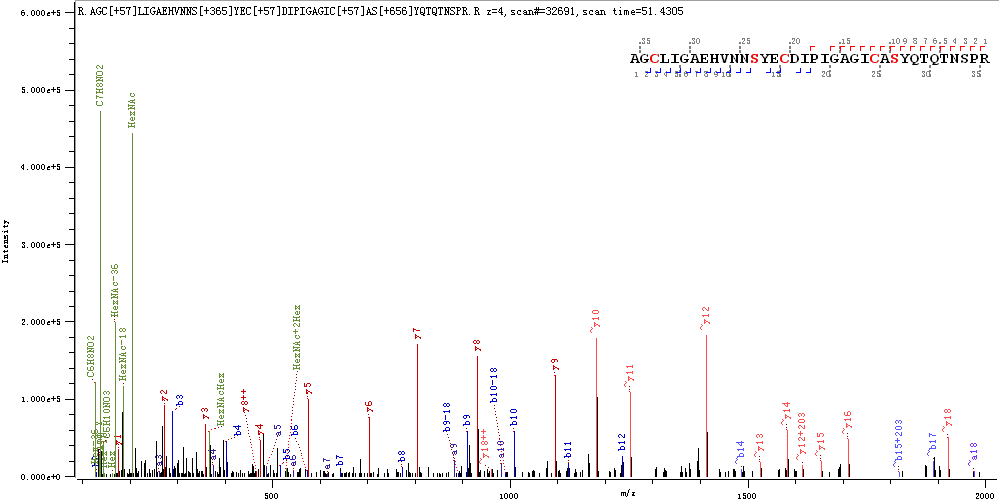

**T732**

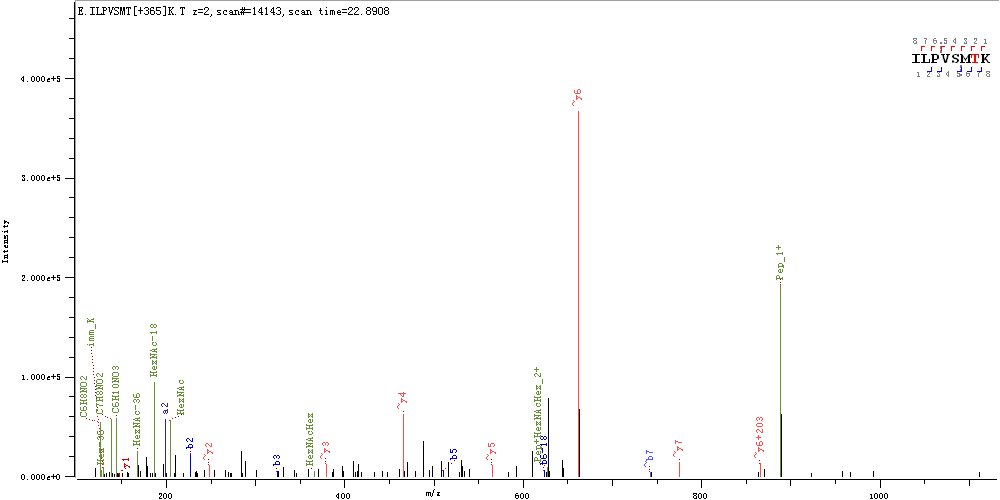

**T791 & S803 & S813**

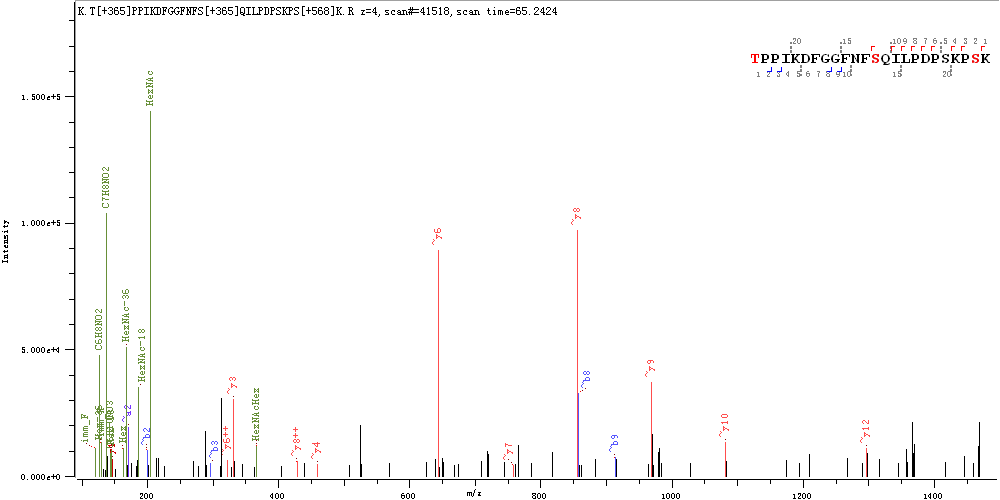

**S810 & S813**

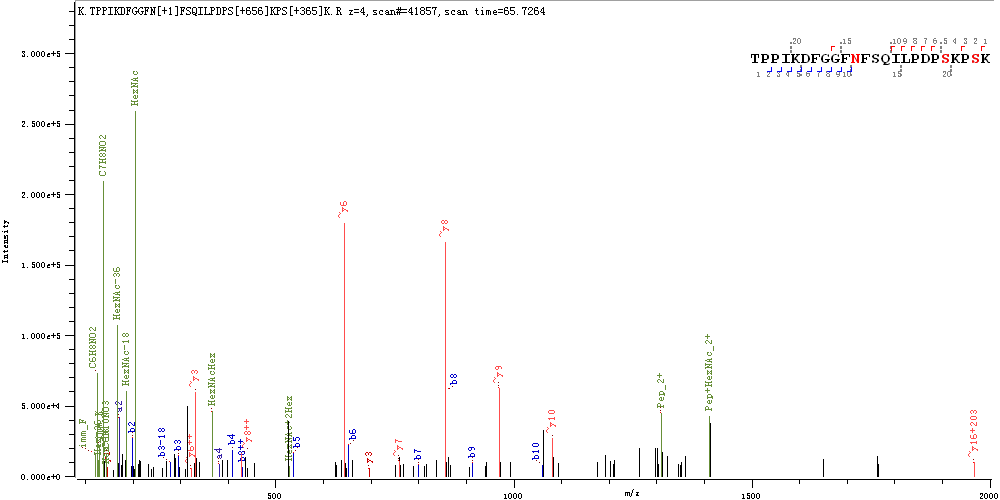

**T912**

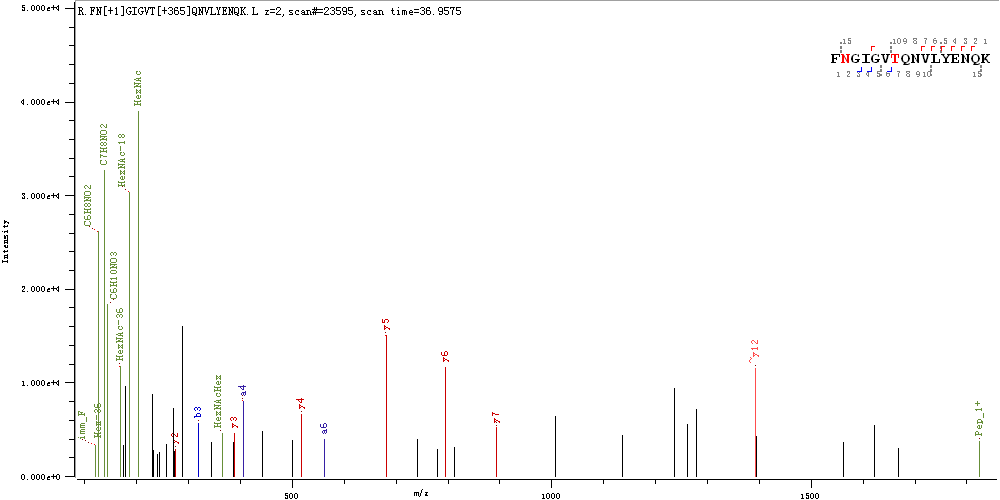

**S939**

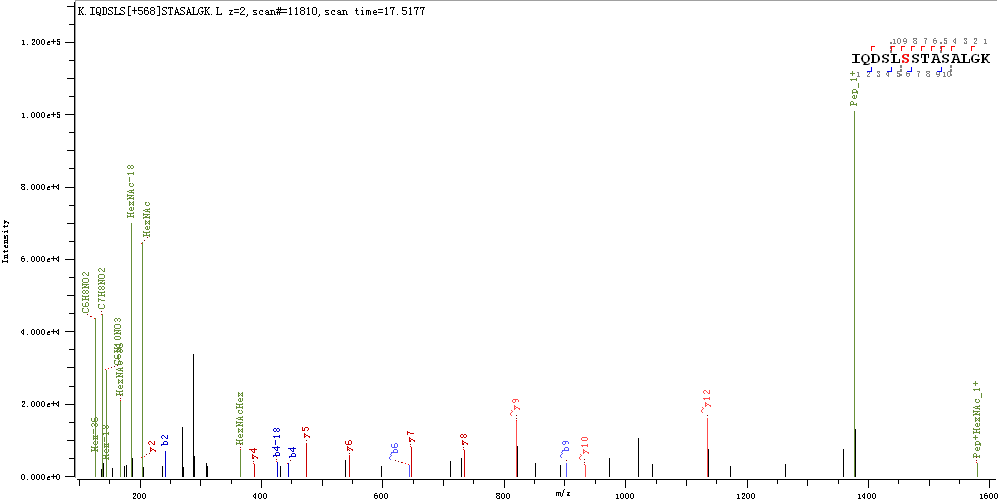

**S940 & T941**

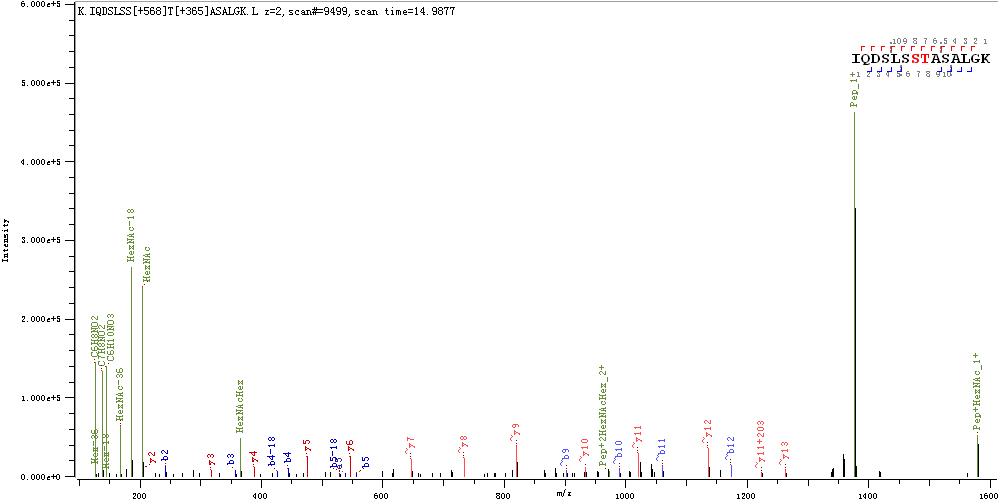

**T1066**

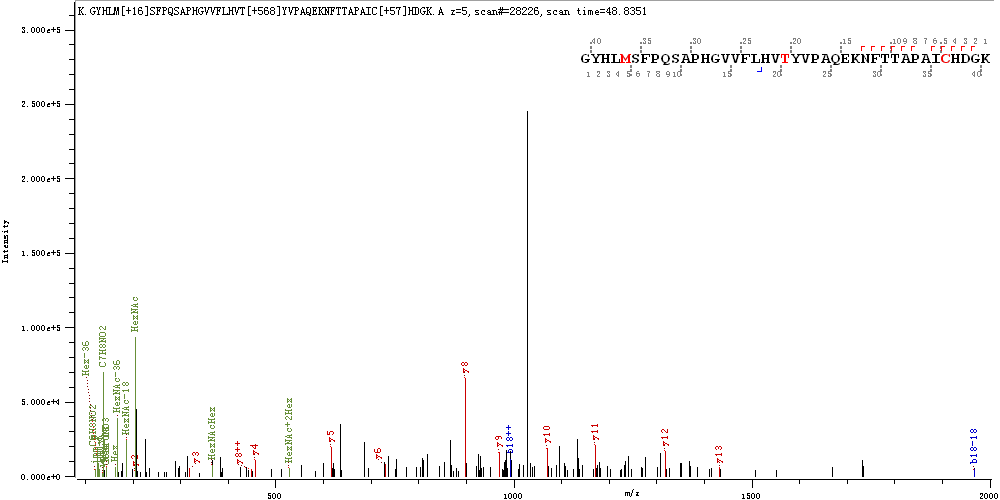

**T1076 & T1077**

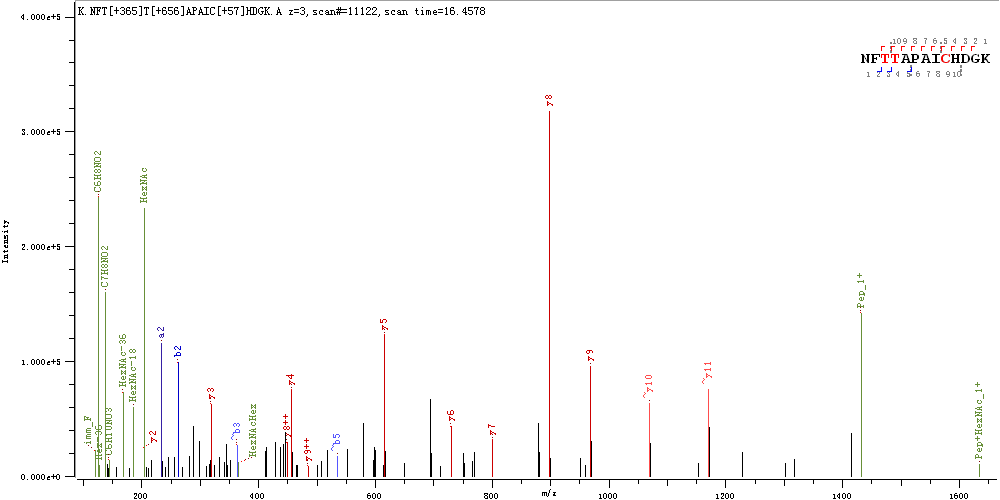

**S1097 & T1100**

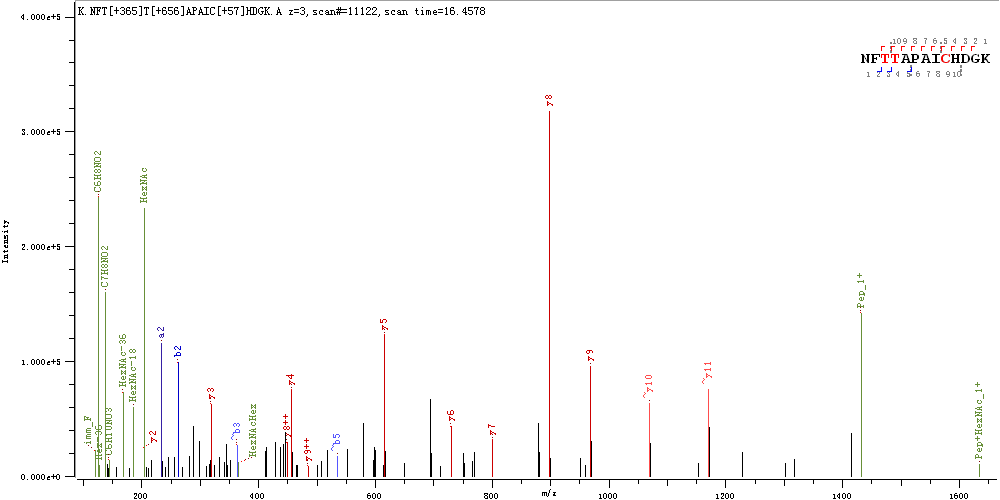

**T1105**

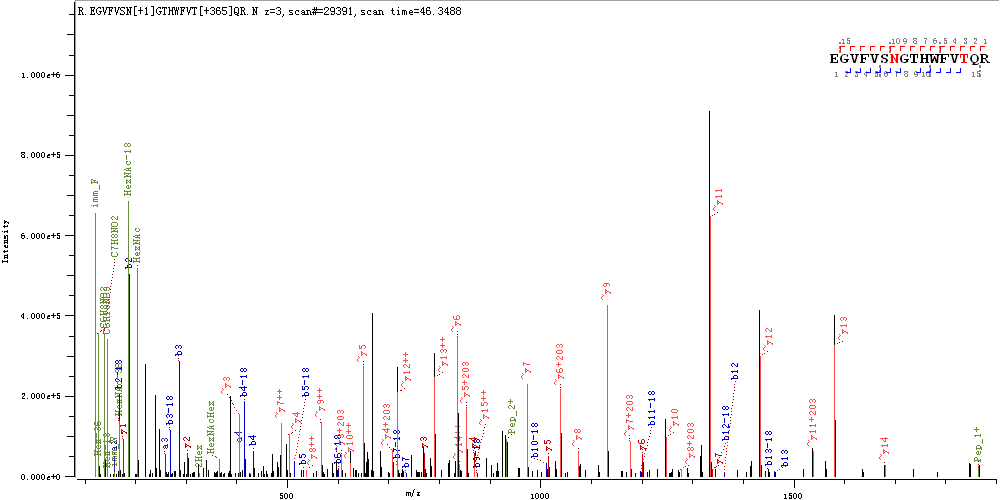

**T1160 & S1170**

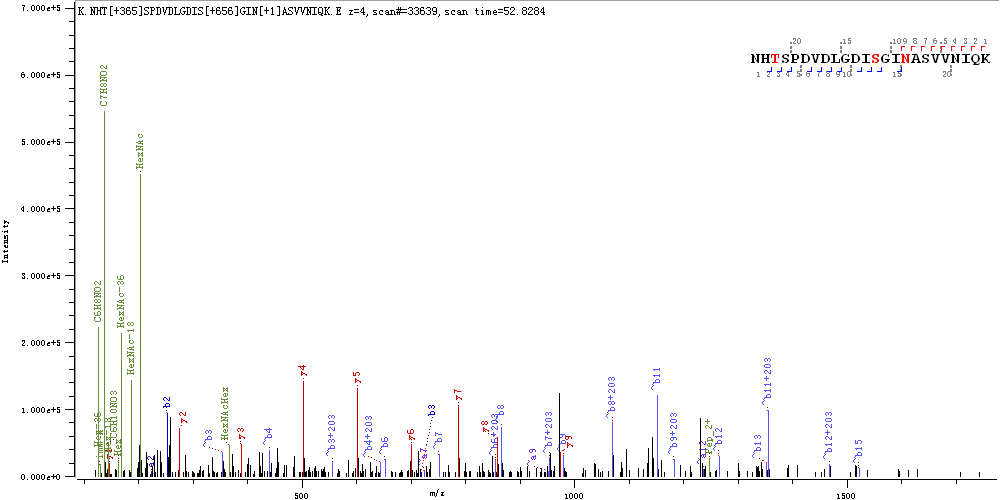

**S1161 & S1170**

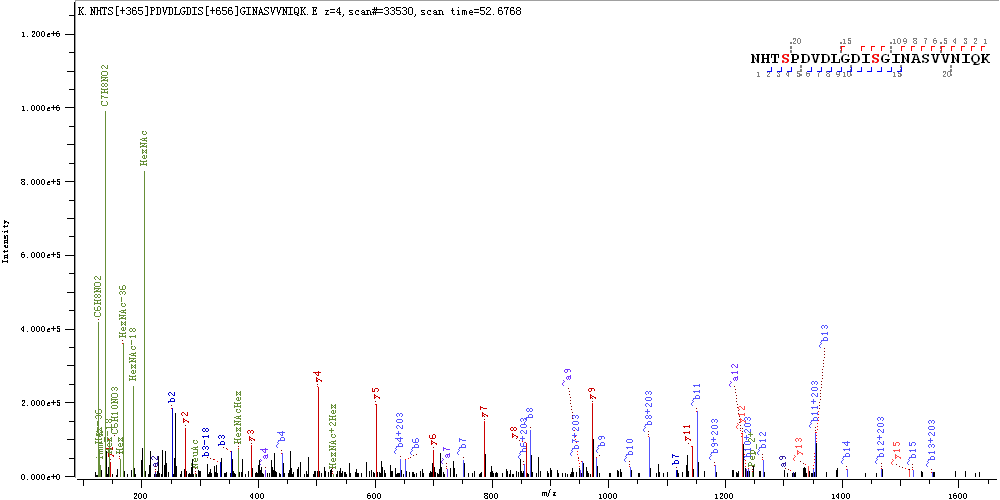

**S1170 & S1175**

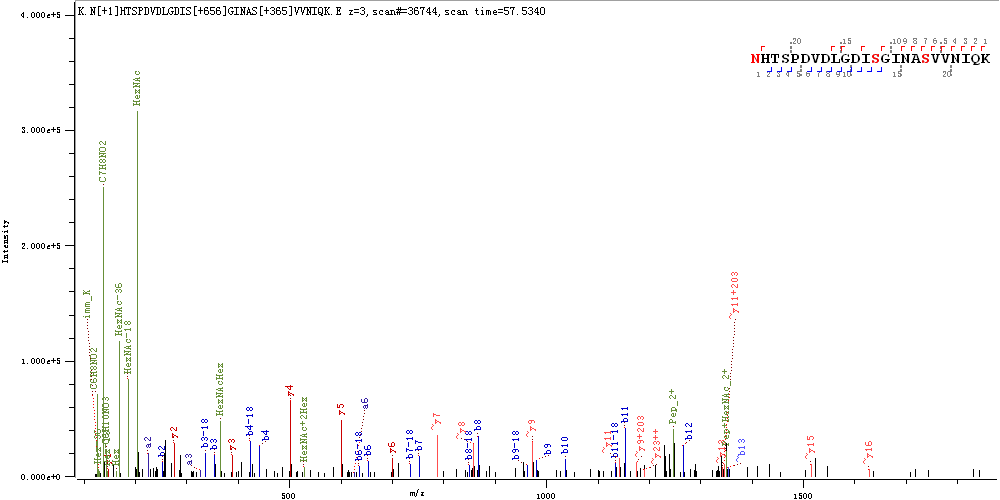

**S1196**

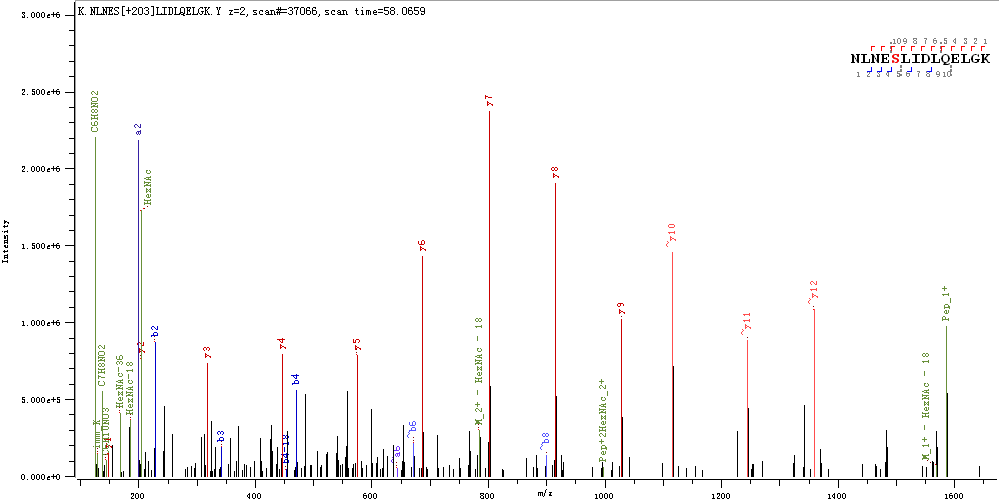

**Supplementary Figure S3.** Spectra of intact *O*-glycopeptides of SARS-CoV-2 S protein expressed in human cells with ambiguously assigned *O*-glycosites

**T22 & T29**

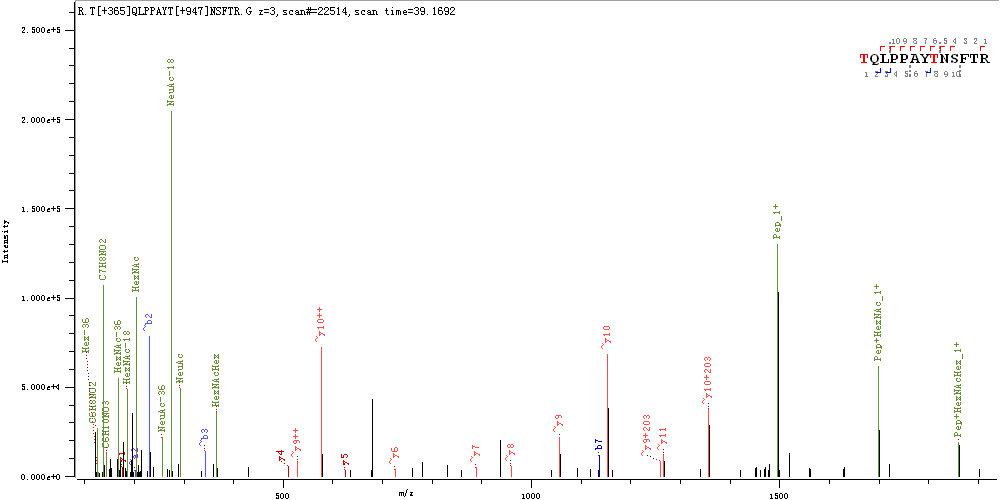

**S31**

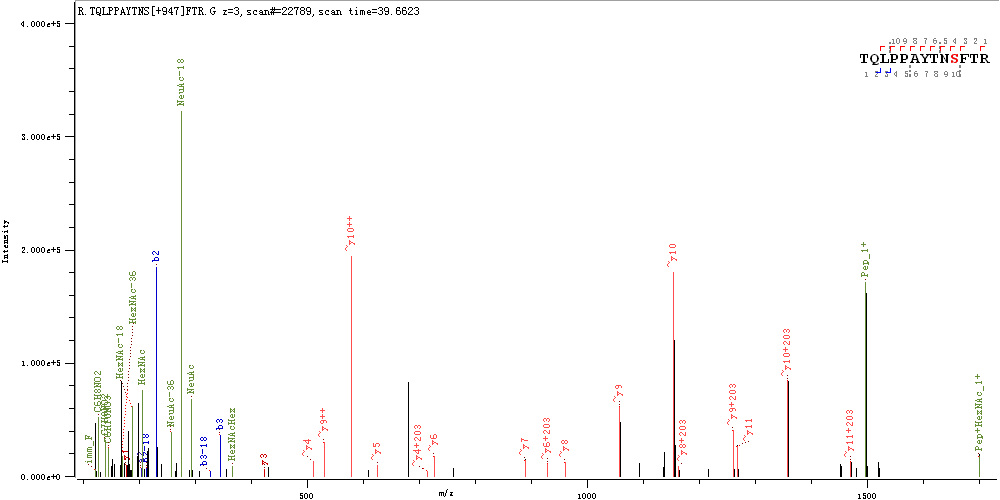

**T33**

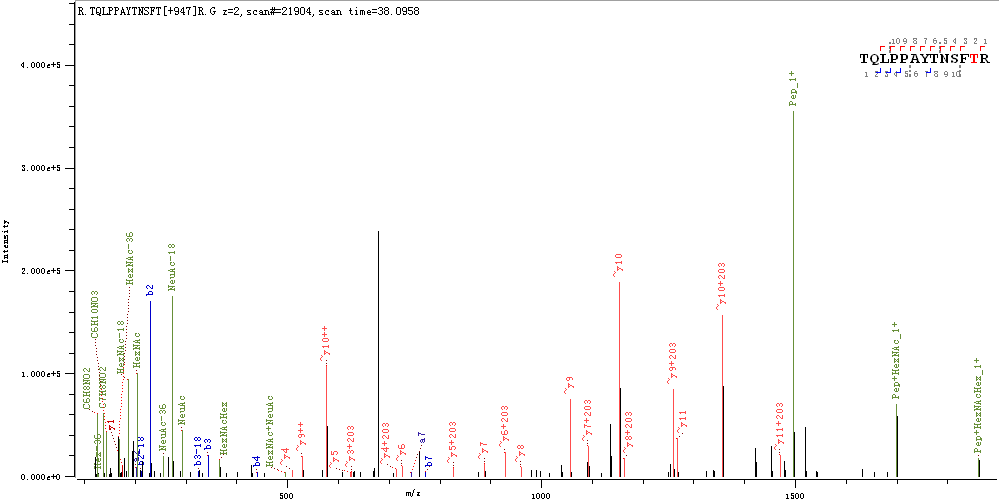

**T124**

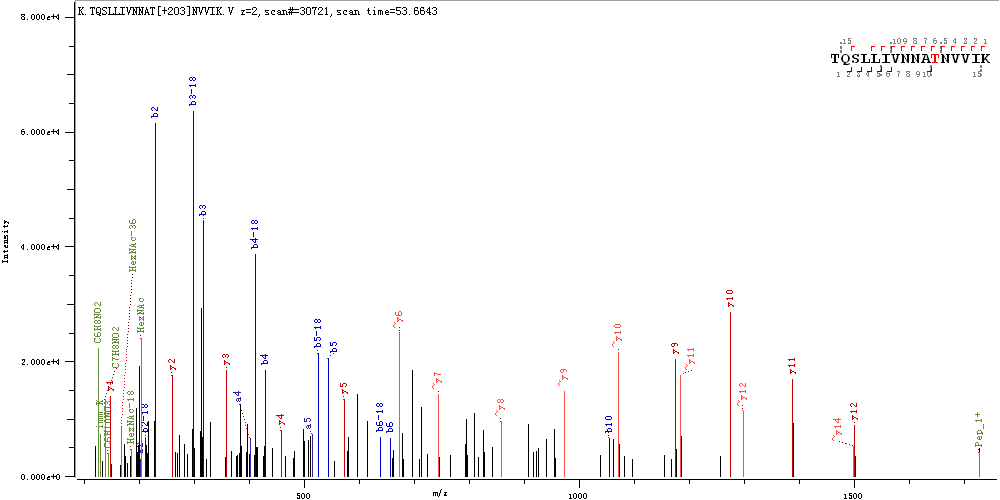

**T284 & T286**

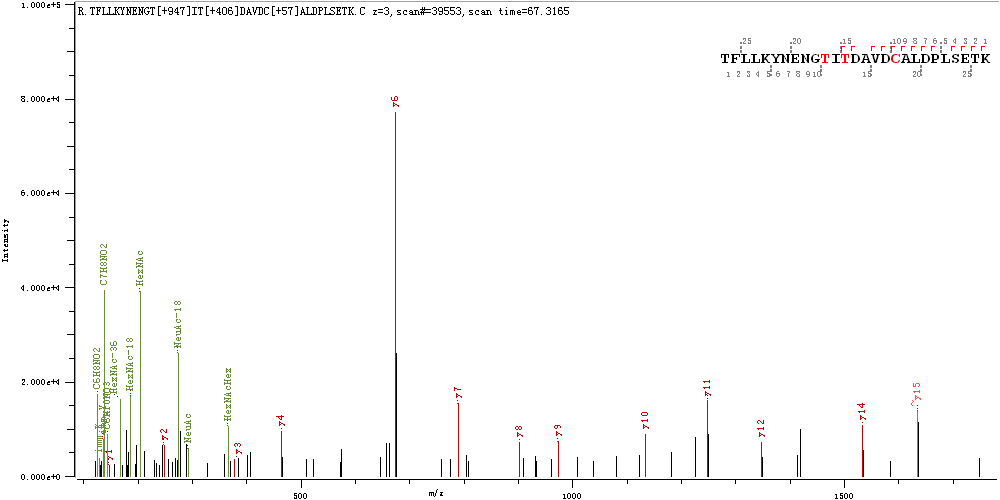

**S297 & T299**

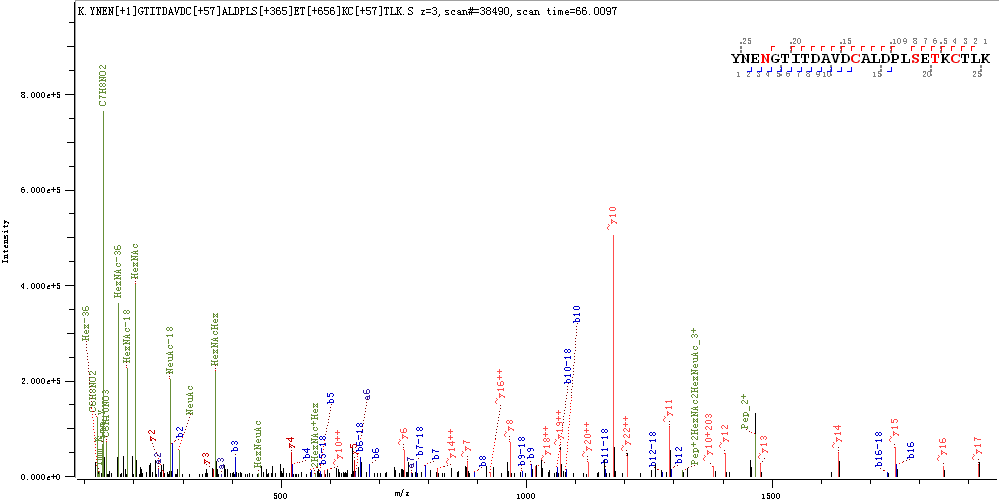

**T302**

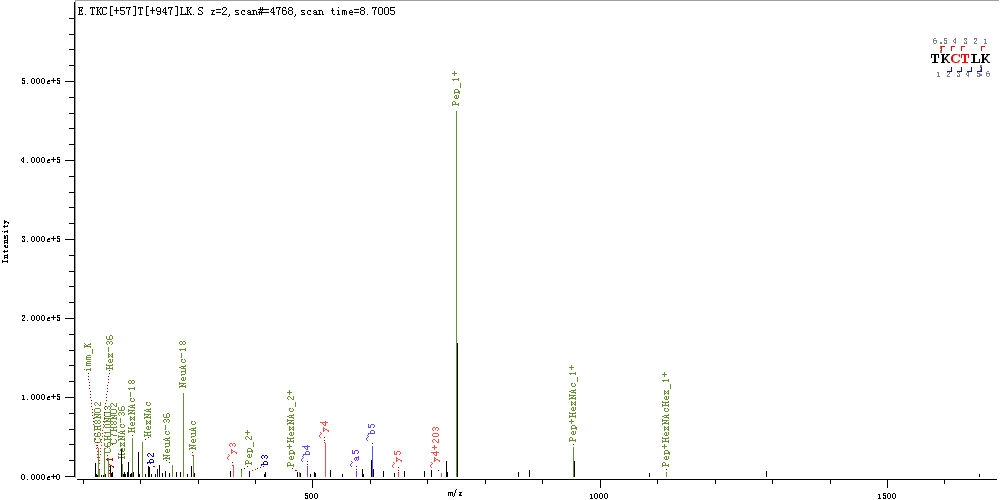

**S305 & T307**

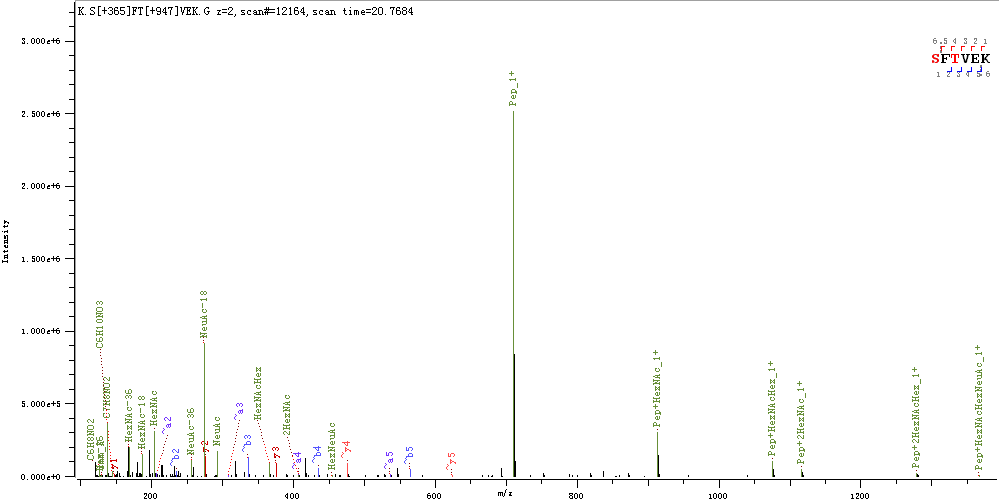

**T315**

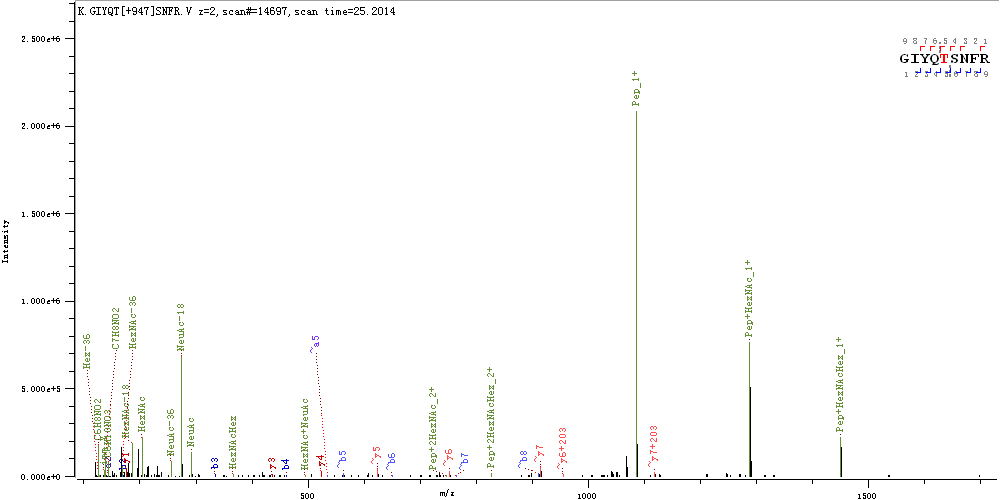

**T315 & S316**

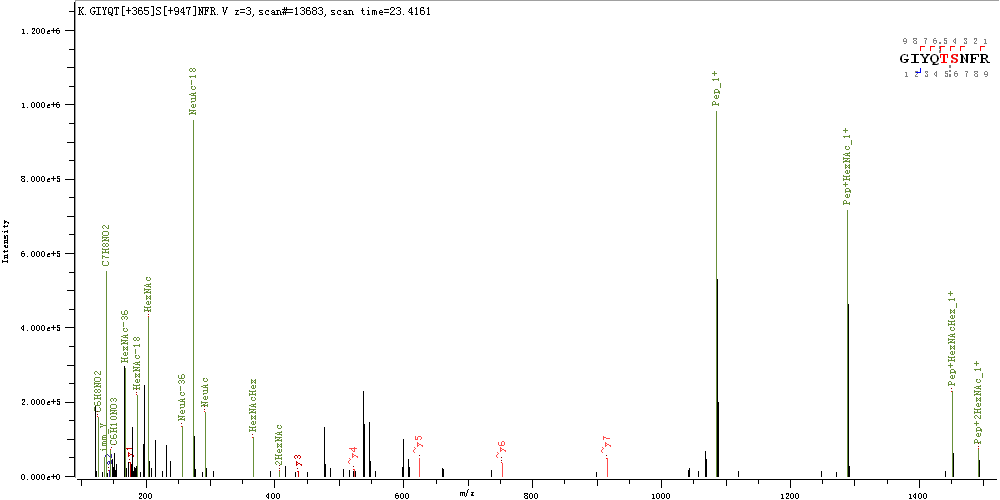

**T323 & S325**

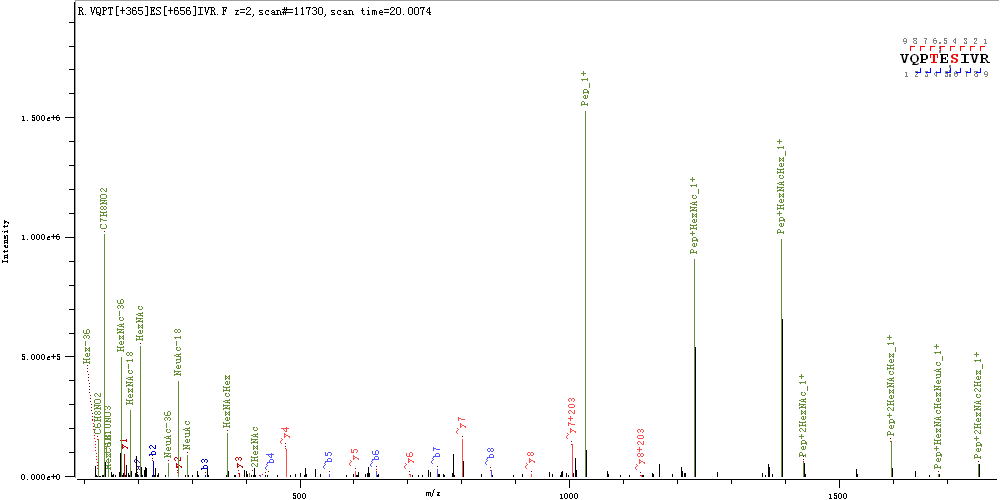

**T572 & T573**

**T573 & T581**

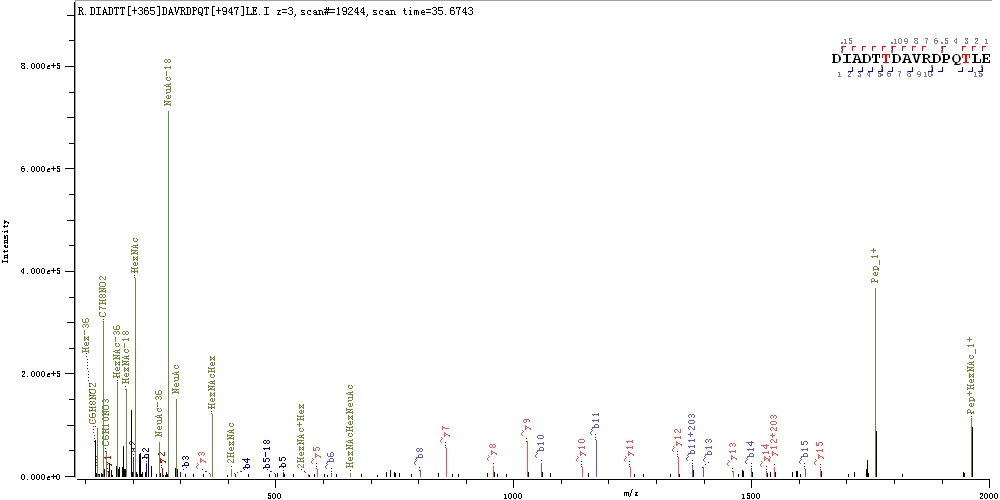

**T630 & T632**

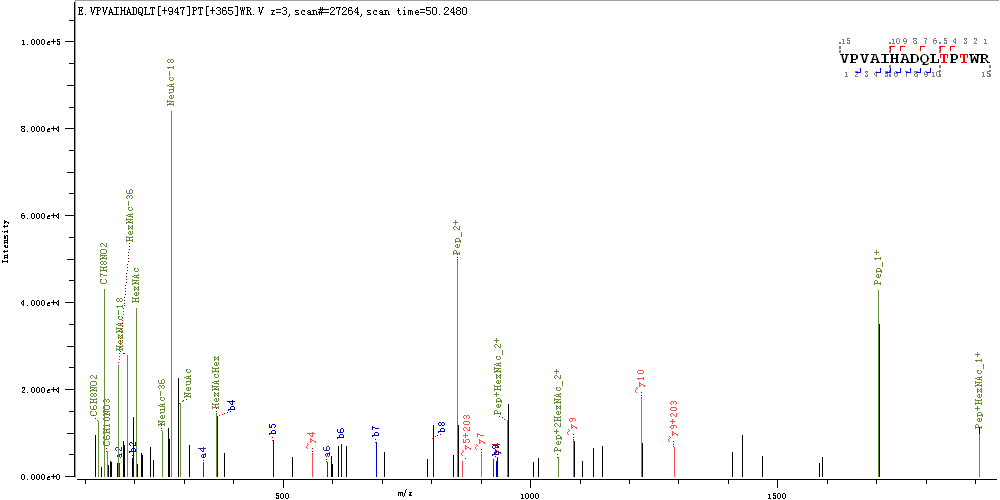

**S637 & T638**

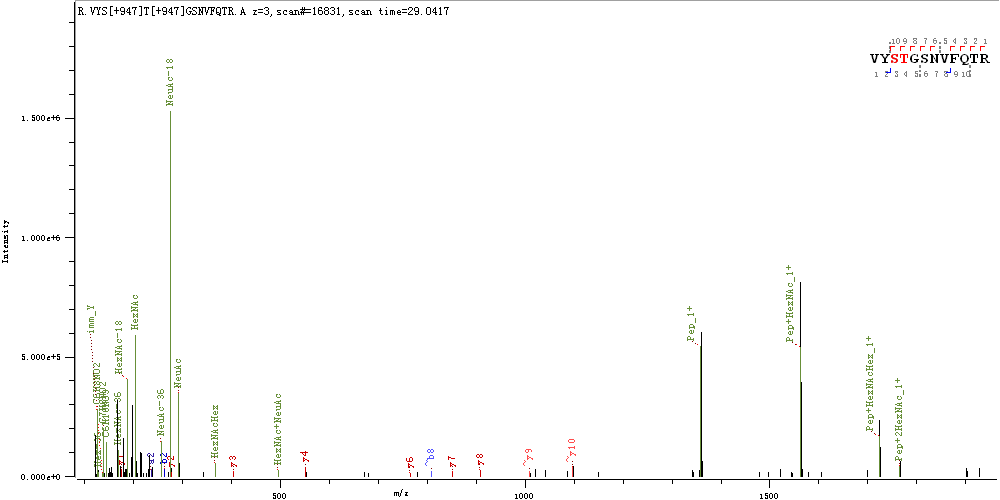

**S640**

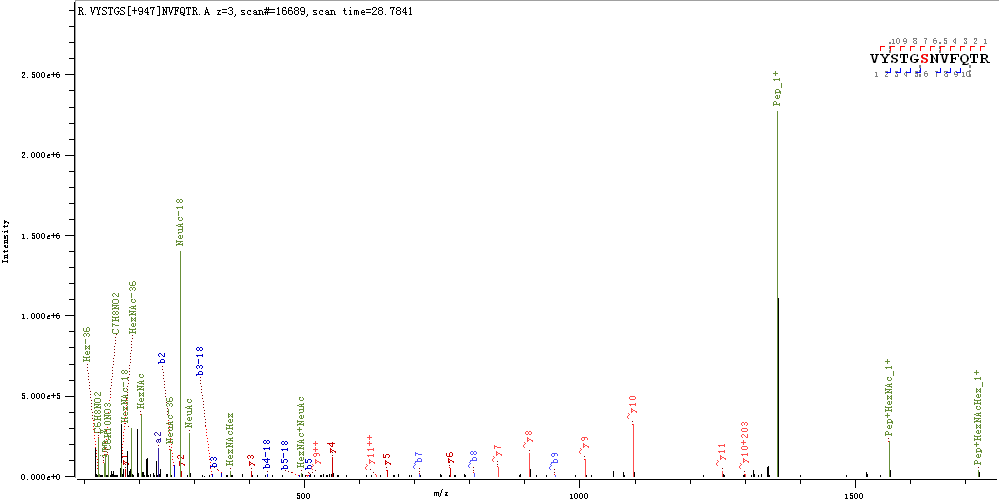

**T645**

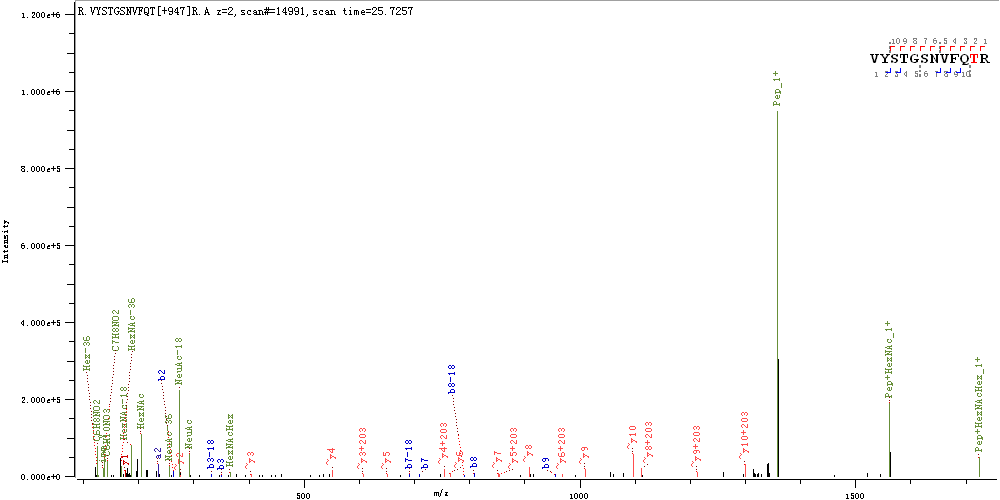

**T659**

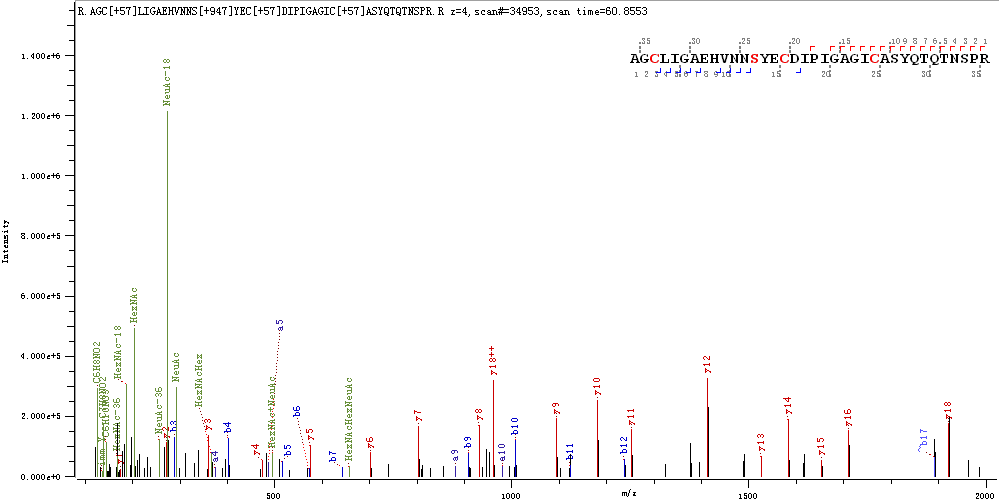

**S673 & S680**

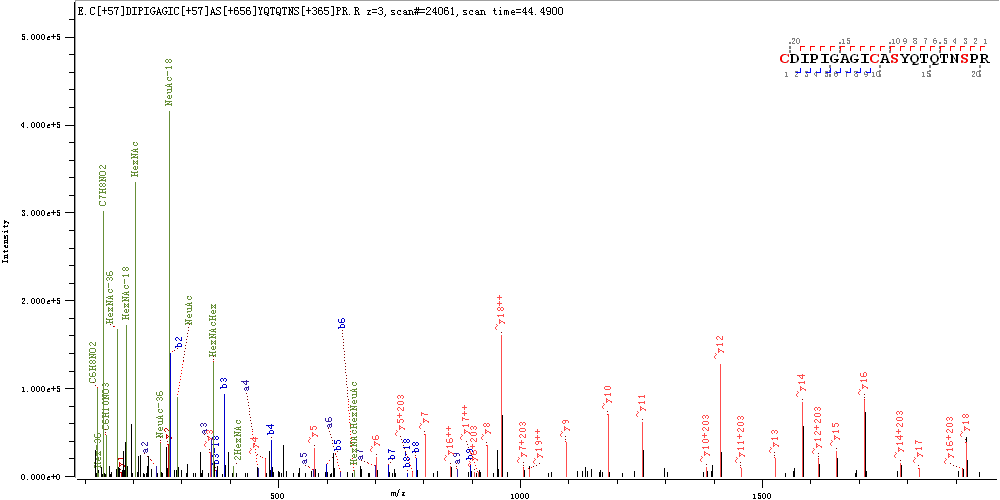

**T676 & S678**

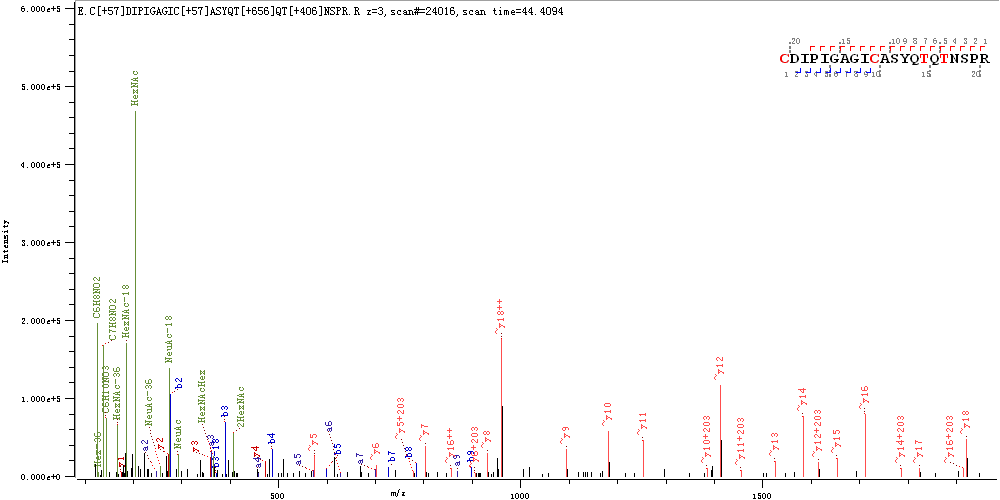
